# Novel peptide targeting CXCR4 disrupt tumor-stroma crosstalk to eliminate migrating cancer stem cells

**DOI:** 10.1101/2025.03.03.641126

**Authors:** Kanishka Tiwary, Anton Lahusen, Syeda Inaas, Bastian Beitzinger, Roman Schmid, Mirja Harms, Stefanie Hauff, Frank Arnold, Karolin Walter, Sonia Alcala, Stephan Hahn, Elisabeth Heßmann, Alexander Kleger, Ninel Azoitei, Thomas Seufferlein, Bruno Sainz, Jan Münch, Mika Lindén, Patrick C. Hermann

## Abstract

Pancreatic ductal adenocarcinoma (PDAC) is one of the most aggressive and metastatic malignancies worldwide. Migrating cancer stem cells (miCSCs) marked by CD133+CXCR4+ expression drives metastasis but lacks effective drug targets. Here, we show that activated pancreatic stellate cells secrete the CXCR4 ligand CXCL12 to foster stemness, epithelial-to-mesenchymal transition (EMT), and chemoresistance. Protein interaction network analyses links CXCL12/CXCR4 signaling axis and the downstream transcription factor BMI1. Knockdown experiments confirmed the BMI1’s role in (mi)CSCs maintenance and survival. Novel CXCR4 inhibitors, i.e., the endogenous human peptide EPI-X4 and its derivatives (e.g., JM#21) strongly inhibited the *in vitro* migration of miCSCs. In particular, the most potent EPI-X4 derivate JM#21 sufficiently suppressed EMT, stemness, and self-renewal of human PDAC cell lines. In addition, JM#21 sensitized cell lines towards gemcitabine and paclitaxel. Overall, our study reveals that (mi)CSCs are enhanced and maintained via a tumor-stroma crosstalk through BMI1, ultimately promoting metastases and therapeutic resistance in PDAC. Peptide targeting of the CXCL12/CXCR4/BMI1 signaling axis via JM#21 could enhance PDAC combination therapies, offering a promising strategy against this deadly cancer.

**Synopsis:** The study identifies a tumor-stroma interaction mediated by pancreatic stellate cells (PSCs) secreting CXCL12, which binds to CXCR4 on (mi)CSCs, fostering stemness, epithelial-to-mesenchymal transition (EMT), and chemoresistance. The CXCL12/CXCR4 axis activates the downstream BMI1 transcription factor, crucial for migration and stemness maintenance.

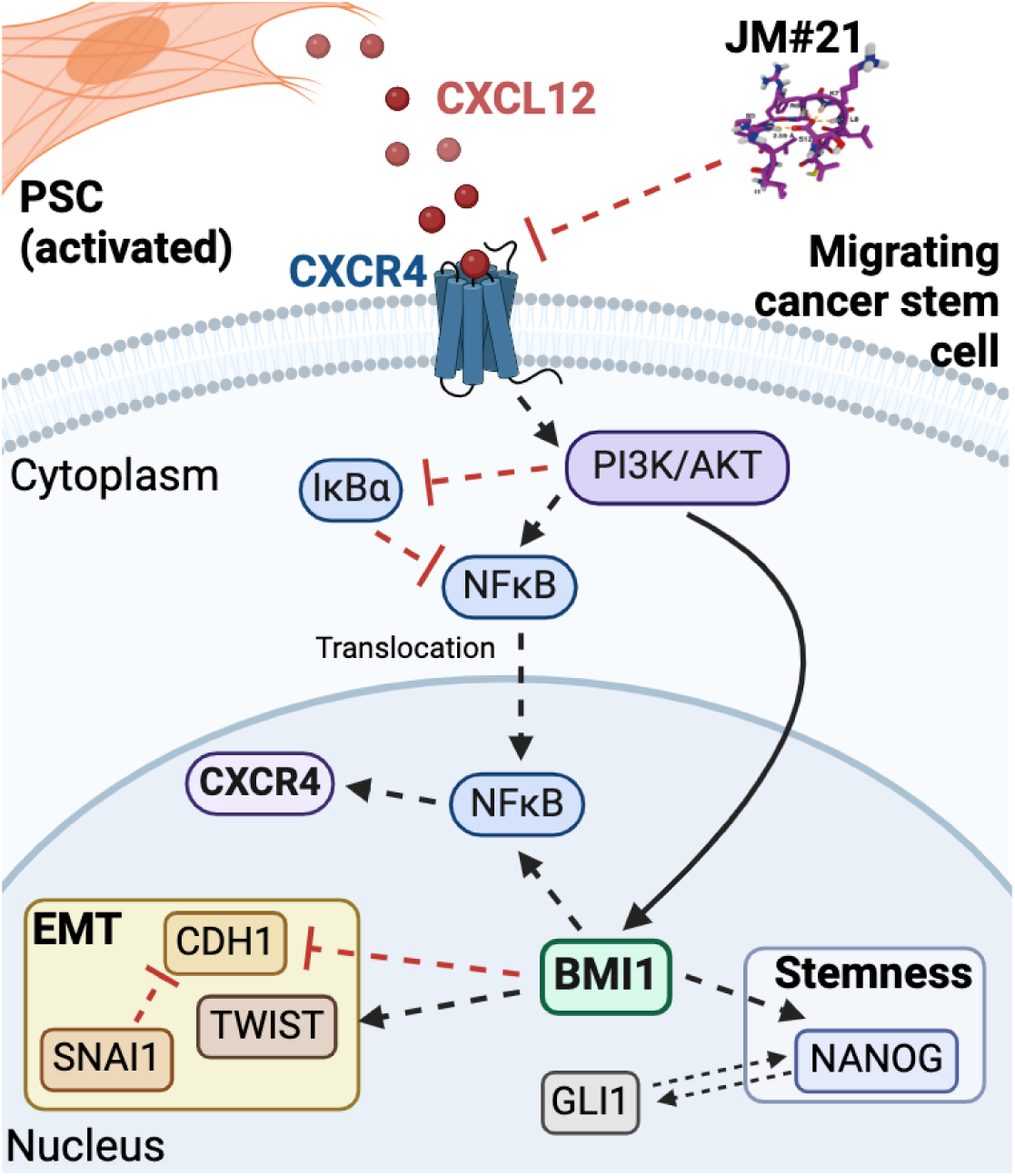

- CXCL12 enhances (mi)CSC populations and metastatic potential through CXCR4 signaling.
- BMI1 is identified as a pivotal downstream effector linking CXCR4 to EMT and stemness.
- JM#21 effectively blocks CXCL12-induced migration, EMT, and stemness in vitro, demonstrating superior efficacy compared to other CXCR4 inhibitors.
- Encapsulation of JM#21 in silica nanoparticles enhances its stability and delivery, reducing chemoresistance and miCSC populations in co-culture systems.
- Combining JM#21 with chemotherapy significantly impairs colony formation and CSC-mediated drug resistance.

## Introduction

Pancreatic cancer accounts for more than a fifth of all gastrointestinal cancer-related deaths and is projected to be the 2^nd^ most frequent cause of cancer-related death by 2030 (Rahib et al., 2014). Histologically, the vast majority of cases (90%) are pancreatic ductal adenocarcinomas (PDAC), which arise from the pancreatic epithelium (Hu et al., 2021). Due to the lack of early symptoms, most PDAC patients present with advanced or metastatic disease at the time of diagnosis (Aier et al., 2019; Sperti et al., 1997), and the 5-year overall survival is only 3% for patients featuring metastasis, an outcome that has largely remained unchanged in the past 15 years (Thomas et al., 2020). For those patients undergoing surgery with a curative intent, nearly 80% ultimately relapse, with 2 out of 3 succumbing to distant recurrence (Sperti et al., 1997).

A reasonable explanation for these alarming statistics is the presence of cancer stem cells (CSCs) within the tumor. CSCs are not only capable of mediating/impacting chemoresistance but also the intrinsic cellular plasticity of tumor cells. At the same time, CSCs can re-populate the tumor via asymmetric and/or symmetric division, even after therapy. Previous studies by Li *et al*. and Hermann *et al*. independently identified CD44+CD24+EpCAM+ cells but also CD133+ cells as CSCs in PDAC, (Hermann et al., 2007; Li et al., 2007). Subsequent studies additionally identified various other markers for pancreatic CSCs (Hermann et al., 2007; Li et al., 2007; Li et al., 2011; Miranda-Lorenzo et al., 2014; Nguyen et al., 2017; Rasheed et al., 2010). Thus, these markers do not identify a pure and exclusive CSC population, but help in advancing our understanding of pancreatic CSC properties, such as their self-renewal capacity, exclusive tumorigenicity or inherent chemoresistance.

We previously identified a subset of CSCs that mediate metastasis, which co-express CD133 and CXCR4 (Hermann et al., 2007). Given their exclusive metastatic activity, they were defined and termed as “migrating CSC (miCSC)”. Interestingly, we documented significantly higher numbers of (CD133+ CXCR4+) miCSCs in patients with lymph node metastases (pN1+), establishing a direct clinical correlation between miCSCs and advanced/metastatic disease. Here, we study the crosstalk between miCSCs and pancreatic stellate cells (PSC), identifying the CXCL12 – CXCR4 axis as being crucial. Specifically, PSC-secreted CXCL12 interacts with CXCR4 to regulate epithelial-to-mesenchymal transition (EMT) and stemness features as well as chemoresistance, orchestrating the phosphorylation of AKT and NFkB and upregulation of BMI1. More importantly, novel endogenous human peptides targeting CXCR4 such as JM#21, an EPI-X4 peptide derivative, resulted in the suppression and subsequent elimination of miCSCs, and can be used without risks of toxicity or off-target effects. To address the translational and potential therapeutic potential of JM#21, we developed a silica nanoparticle (SiNP)-encapsulated JM#21 formulation which substantially improves stability and delivery of the compound. Of note, SiNP-encapsulated JM#21 drastically reduced chemoresistance and stemness features in tumor cell – stellate cell co-culture systems, supporting the potential of peptide CXCR4 inhibitors as a putative therapeutic avenue to eliminate miCSCs and metastasis in PDAC.

## Results

### CD133 and CXCR4 are overexpressed in PDAC

In previous studies interrogating the role of CXCR4 in PDAC and employing its pharmacologic abrogation with the CXCR4 inhibitor AMD3100, we identified CD133+ CXCR4+ cells as migrating CSCs (Hermann et al., 2007). In order to understand the intimate mechanisms by which CD133 and CXCR4 may contribute to CSC features and EMT, we performed *in-silico* analysis of the Cancer Genome Atlas (TCGA) and GTEx PDAC database. Differential expression analysis indicated significantly elevated levels of *PROM1* (CD133) and/or *CXCR4* in PDAC as compared to normal tissue (**Fig. 1A**). At the same time, a STRING network analysis revealed key interactions of CD133 and CXCR4 with other relevant factors involved in metastasis and EMT (green) or CSCs (red) (**Fig. 1B**). To further substantiate these findings, we investigated the levels of CD133 and CXCR4 by employing 9 primary cell lines derived from patient-derived primary tumor xenografts (PDX) and patient-derived organoids (PDO). Flow cytometry analysis revealed varying levels of CD133+ CSC (**Fig. 1C**) and CD133+ CXCR4+ miCSC (**Fig. 1D**) populations in the investigated cell lines.

**Figure 1.**
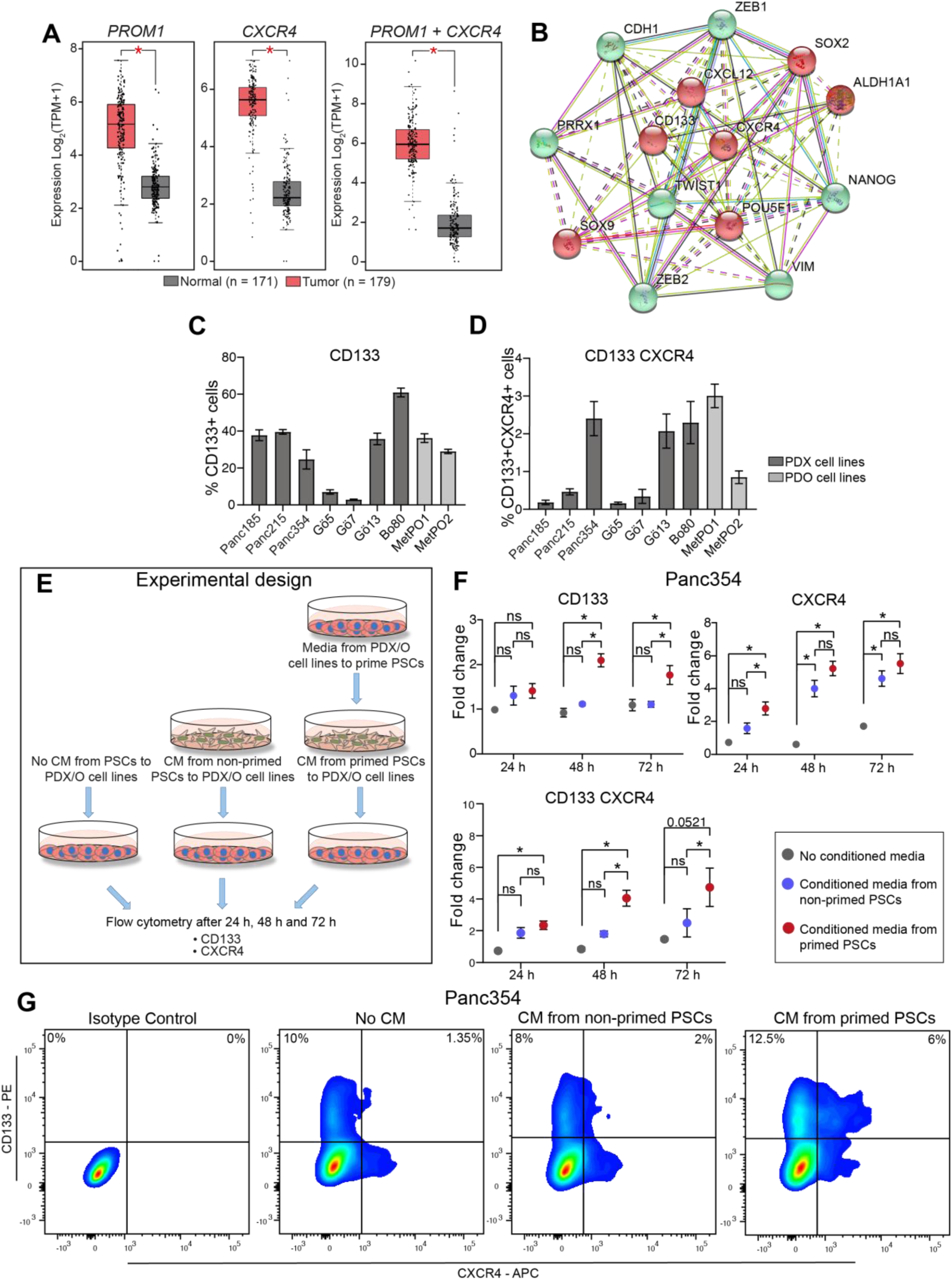
CD133 and CXCR4 are overexpressed in PDAC. (A) Box plots represent relative mRNA expression for the gene signatures *PROM1, CXCR4 or PROM1 and CXCR4* in normal tissue compared to PDAC tumor tissue (TCGA and GTEx database, GEPIA). (B) Protein – protein interactions of PROM1 (CD133) and CXCR4 with relevant factors involved in metastasis (green) and CSCs (red) (STRING). (C) Screening for percent CD133+ cells and (D) CD133+ CXCR4 + found in adherent cell culture of indicated cell lines. (E) Experimental scheme to evaluate CD133 and CXCR4 surface expression to identify patient derived xenografts (PDX) or patient derived organoids (PDO) crosstalk with pancreatic stellate cells (PSCs) at the protein level. (F) FACS analysis performed on Panc354 and MetPO1 when exposed to no conditioned media (grey), conditioned media from non - primed PSCs (blue) or conditioned media from primed PSCs (pink) for CD133+ cells, CXCR4+ cells and CD133+ CXCR4+ cells represented as fold change against no conditioned media. (G) Representative cytometry blots. Error bars represent the standard deviation. n≥3 for all experiments. *p < 0.05, ns = not significant.

There is intense crosstalk between pancreatic CSCs and the microenvironment (Ligorio et al., 2019). Pancreatic stellate cells (PSC) are part of the supportive niche for pancreatic CSCs and secrete a plethora of factors that are crucial for promoting PDAC heterogeneity (Ligorio et al., 2019). Therefore, we sought to evaluate the contribution of these factors for the maintenance of CSC and miCSC. We speculated that CSCs and miCSCs within its microenvironment secretes factors that might educate or prime PSCs to their presence. To capture that, conditioned media (CM) was prepared using media from PDX-derived Panc354 and PDO-derived MetPO1 cell lines and was used to tumor “educate” or “prime” the PSCs (**Fig. 1E**). Next, Panc354 and MetPO1 cell lines were exposed to conditioned media (CM) of non-primed PSC, primed PSC as well as control media (**Fig. 1E**). Interestingly, cultivation with CM harvested from primed PSCs (red) were associated with higher levels of CD133+, CXCR4+ and CD133+CXCR4+ cells (**Fig. 1F-G; Supp. Fig. 1A**) when compared to CM from non-primed PSCs (blue) or no CM (gray). Our findings suggest that while PSC-secreted factors indeed upregulate CD133 and CXCR4, conditioned media from PDX- and PDO-primed PSCs further boost the levels of CD133 and CXCR4 in pancreatic cancer cells, suggesting that the secretome of activated PSCs can further increase the potency of CSCs and miCSCs (**Fig. 1F-G; Supp. Fig. 1A**).

### Tumor cell – stellate cell crosstalk promotes CSC and miCSC populations and potentiates CXCL12 release

The communication between tumor cells and stellate cells is of paramount importance for maintaining the traits of CSCs and miCSCs. In addition to the experiments using conditioned media, PDX/PDO cell lines were co-cultured with pancreatic stellate cells (or PDX/PDO cell lines as control) using culture inserts for 72 hours. Cells were harvested and subjected to further analysis as depicted in (**Fig. 2A**). Flow cytometry analysis again revealed a significant increase in CD133 and CXCR4 in both CSCs and miCSCs (**Fig. 2B-D**). Gene expression analysis from PDX/PDO derived cell lines co-cultured with PSCs revealed significant upregulation of stemness-associated genes (*NANOG, POU5F1, ALDH1a1, BMI1*) (**Fig. 2E**). Interestingly, *DCK1*, a gene responsible for catalyzing the rate-limiting step in gemcitabine metabolism (Hu et al., 2018) was downregulated in our experimental setting (**Fig. 2E**). Moreover, *ABCC5*, a gene involved in the transport of nucleotide analogues, and previously speculated to be responsible for the excessive efflux of nucleotide analogue-based drugs such as 5-FU or gemcitabine (Hagmann et al., 2010), was suppressed in all tested cell lines. At the same time, we observed regulation of genes involved in the metabolism of reactive oxygen species (ROS) (Cheung et al., 2020; Zhang et al., 2016): while *SOD1* was upregulated, *GPX1* displayed a rather heterogeneous expression across PCSs with elevated expression restricted to Panc354 – PSCs only. Importantly, the epithelial marker *CDH1* was significantly downregulated, while EMT-specific markers (*VIM, SNAI1, SNAI2*) showed elevated expression levels (**Fig. 2E**), indicating a more mesenchymal phenotype. Altogether, these findings indicate a strong reshaping of the transcriptional profile of PDX/PDO cell lines during co-culture with PSCs. These findings were further substantiated by functional assays: PDX/PDO-derived cell lines (either co-cultured with PSCs or PDX/PDO) were analyzed in 3D transwell migration assays (**Fig. 2F-G**) and sphere formation assays (**Fig. 2H-I**), indicating that PSCs significantly potentiated the migratory capability and sphere formation potential of PDX/PDO-derived cell lines.

**Figure 2.**
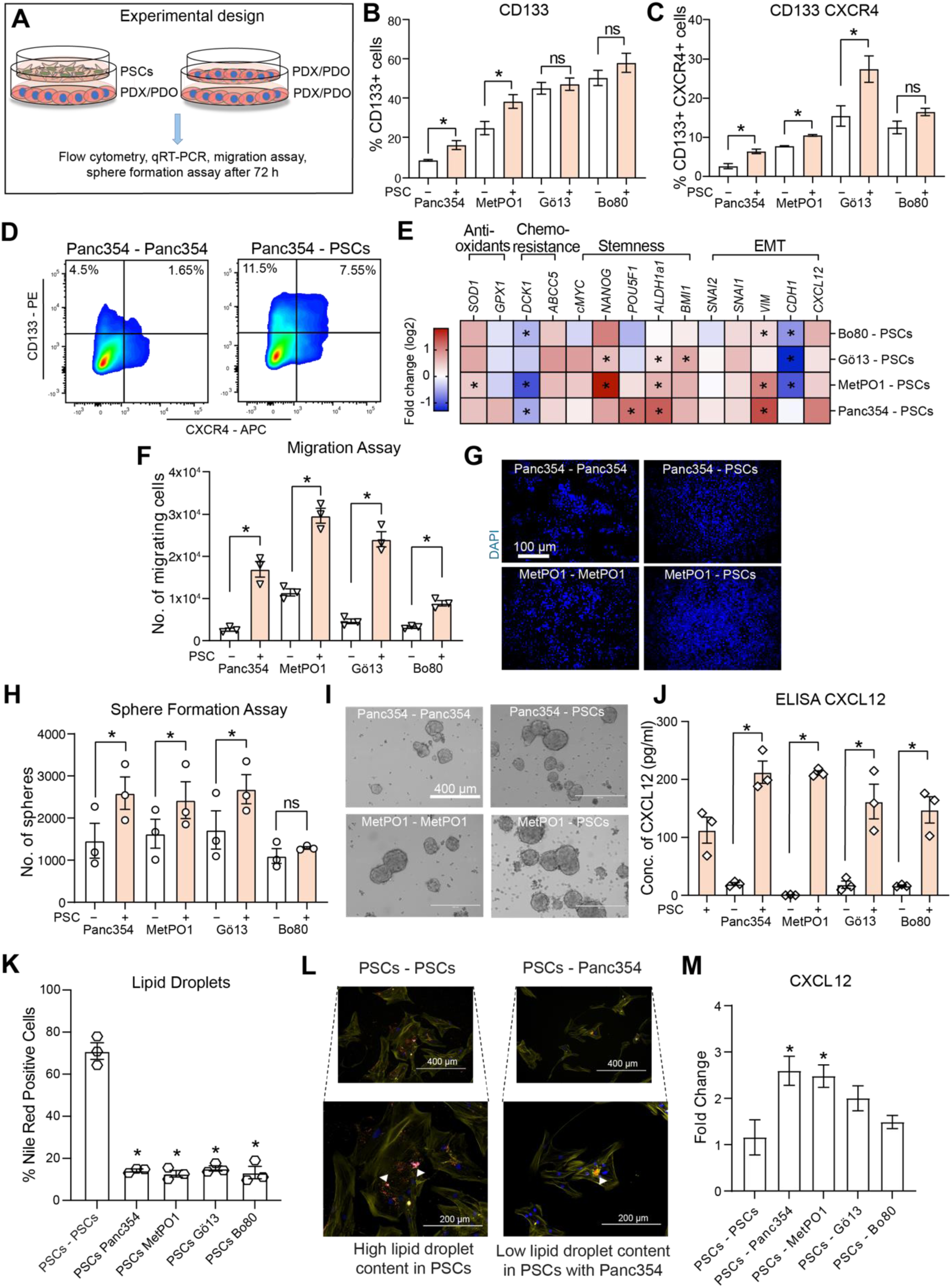
Tumor cell – stellate cell crosstalk upregulates CSC and miCSC population, and potentiates CXCL12 release. (A) Experimental scheme to evaluate CD133 and CXCR4 surface expression, qRT-PCR, migration assay, sphere assay, ELISA and IF for co-culture PDX/PDO – PSCs and PDX/PDO – PDX/PDO. (B) Flow cytometry analysis performed for percent CD133+ cells and (C) CD133+ CXCR4+ cells for depicted cell lines. (D) Representative cytometry plots. (E) Gene expression analysis for indicated genes using qRT-PCR for total RNA extracted from PDX/PDO cell lines (indicated) co-cultured with PSCs or PDX/PDO cell lines represented as log2 fold change. (F) Transwell migration assays for PDX/PDO cell lines (indicated) cultivated in dual culture with PSCs or PDX/PDO cell lines and (G) representative micrographs (10x, DAPI nuclear staining). (H) Sphere formation assays for PDX/PDO cell lines (indicated) cultivated in dual culture with PSCs or PDX/PDO cell lines and (I) representative micrographs of sphere cultures after 7 days. (J) ELISA performed for CXCL12 concentration (pg/ml) in media from cultures as indicated. (K) Percentage of Nile red (lipid droplet) positive PSCs grown in 2D cultures with different cell lines as indicated and (L) representative micrographs with white arrowheads marking lipid droplets stained with Nile Red (red dots), nucleus with DAPI (blue) and actin filaments with Phalloidin (yellow). (M) Gene expression analysis for CXCL12 using qRT-PCR for total RNA extracted from PSCs cell lines (indicated) co-cultured with PSCs or PDX/PDO cell lines represented as fold change. Error bars represent the standard deviation. n≥3 for all experiments. *p < 0.05, ns = not significant.

CXCL12 was previously reported to be abundantly expressed in the TME (Pan et al., 2015) and typical target organs of metastasis such as liver, lung, lymph nodes etc. (Muller et al., 2001), and to bind to CXCR4 (Bleul et al., 1996; Lopez-Gil et al., 2021; Oberlin et al., 1996). Furthermore, we previously showed in a study on CSC-dependent migration and metastasis of PDAC that metastatic spread is dependent on both CXCR4 expression on tumor cells and the amount of CXCL12 in the target organ (Muller et al., 2001). In order to elucidate whether CXCL12 contributed to the migratory events observed in our experimental setting, CXCL12 concentrations in media from PSCs alone and PSC-PDX/PDO co-cultures was measured by ELISA (**Fig. 2J**). While PSC monocultures were associated with a significantly higher amount of CXCL12 when compared to PDX/PDO cultures, co-cultivation revealed a boost in CXCL12 secretion, indicating the activated (or primed) state of PSCs (Pan et al., 2015).

PSC activation upon co-culturing PSCs with PDX/PDOs was validated by decreased lipid droplet content, assessed by Nile Red staining (**Fig. 2K-L**). These findings correlated with a strong expression of *CXCL12* within the PSCs upon co-culture with PDX/PDO cells lines (**Fig. 2M**). Altogether, these findings indicate an intimate communication between tumor cells and PSCs that ultimately translates into an augmented self-renewal capacity and invasiveness of tumor cells and a boost in the release of CXCL12 by stellate cells.

### CXCL12 – CXCR4 axis sustains the self-renewal capacity and metastatic propensity of miCSCs

Having shown both an increase in CXCL12 and CXCR4 in our co-culture model, we next investigated whether the cytokine funnels its signals directly through CXCR4. We chose to proceed further with two primary cultures that expressed comparable levels of CD133 and CD133/CXCR4, but that have different metastatic capacities: 354 (non-metastatic) and MetPO1 (metastatic) (**Fig. 1C-D**). sh-RNA mediated knockdown of CXCR4 was successfully achieved in both cell lines (**Fig. 3A; Supp. Fig. 1B-C**). MetPO1 cells with CXCR4 knockdown (KD) were then cultured in the presence or absence of CXCL12 for 24 hours. In parallel, CXCR4-KD MetPO1 cells were co-cultured with PSCs for 72 hours and then subjected to further experiments as illustrated in **Fig. 3B**. In sh_SCR cells, both CXCL12 treatment or the co-cultivation with PSCs resulted in a substantial increase in miCSCs (CD133+CXCR4+) cells. Interestingly, CXCR4 KD alone was associated with a decrease in CD133+ cells and CD133+CXCR4+ cells (**Fig. 3C-E**), and exposure to CXCL12 or co-cultivation with PSCs did not restore CD133+ or CD133+CXCR4+ expression to the respective shRNA control levels after CXCR4 abrogation in Panc345 and MetPO1 cells (**Fig. 3C-E; Supp. Fig. 1D**). These findings support a critical role for the CXCR4-CXCL12 axis in CSC propagation. In line with these findings, CXCR4 deletion also decreased sphere formation capacity and migratory potential promoted by CXCL12 as shown in **Fig. 3F-G (Supp. Fig. 1F-G**). Furthermore, gene expression and protein analyses revealed the upregulation of several EMT factors and cancer stemness markers upon incubation with CXCL12 or co-cultivation with PSCs (F**ig. 3H-I** ; **Supp. Fig. 1E and 1H**). The potential of CXCL12 and/or co-cultivation with PSCs to activate EMT and stemness factors was substantially suppressed after CXCR4 abrogation in MetPO1 cells (**Fig. 3I**). Our investigations also demonstrate that genes involved in imparting/rendering chemoresistance in PDAC, such as *CDA1* (Bjanes et al., 2020) and *ABCC5* (Chen et al., 2021) were significantly overexpressed upon treatment of control cells with CXCL12 or co-cultivation with PSCs **(Fig. 3H)**. This was further associated with significant upregulation of antioxidant enzymes such as *GPX1* and *SOD1,* which are responsible for catalytic transformation of reactive oxygen species and their by-products into stable nontoxic molecules (Mates et al., 1999). Abrogation of CXCR4 suppressed chemoresistance and antioxidant enzyme genes induced by CXCL12- and / or PSCs **(Fig. 3H; Supp. Fig. 1E)**. Finally, loss of CXCR4 inhibited CXCL12-induced phosphorylation of AKT and IκB-α and thus NFκB pathway activity, further affecting cell survival **(Fig. 3J; Supp. Fig. 1G)**.

**Figure 3.**
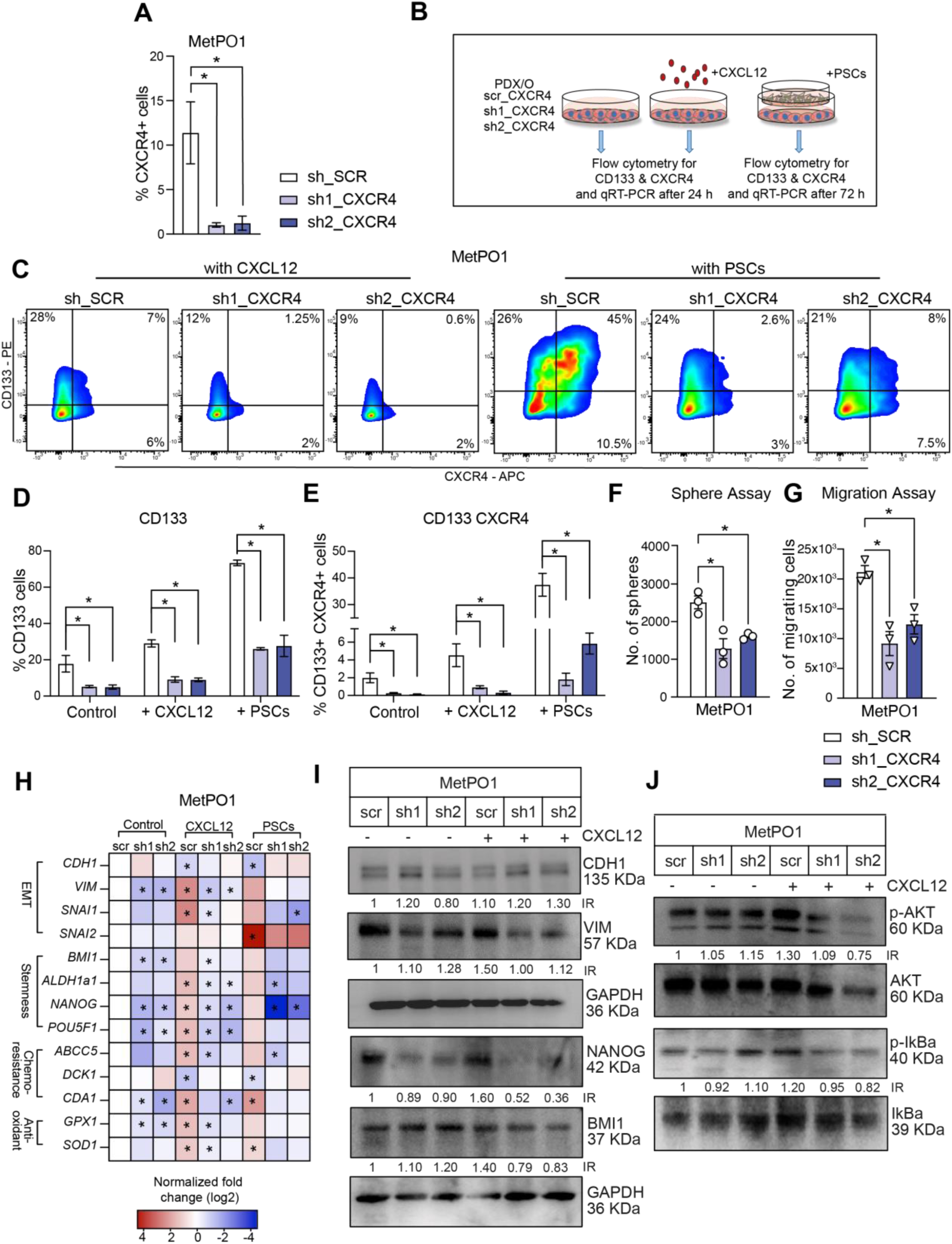
CXCL12 – CXCR4 axis sustains the self-renewal capacity and metastatic propensity of miCSCs. (A) Flow cytometry analysis of CXCR4 surface expression in depicted cell lines. (B) Experimental setup to evaluate CD133 and CXCR4 surface expression and qRT-PCR for Panc354 and MetPO1 (sh_*SCR*, sh1_*CXCR4* and sh2_*CXCR4*). (C) Representative cytometry plots for MetPO1 cell line for flow cytometry analysis of (D) CD133+ CSCs and (E) CD133+ CXCR4+ miCSCs in MetPO1 (sh_*SCR*, sh1_*CXCR4* and sh2_*CXCR4*) treated with (or without) CXCL12 or co-cultured with PSCs. (F) Sphere formation assay after CXCL12 treatment. (G) Migration assay towards CXCL12. (H) Targeted gene expression analysis for indicated genes in MetPO1 (sh_*SCR*, sh1_*CXCR4* and sh2_*CXCR4*) treated with (or without) CXCL12 or co- culture with PSCs. (I) Western blot analysis of labelled protein markers for MetPO1 cell line and treatment conditions. GAPDH was used as a loading control. Error bars represent the standard deviation. n≥3 for all experiments. *p < 0.05, ns = not significant.

### BMI1 regulates EMT and stemness through the CXCL12 – CXCR4 downstream effector axis

To delineate critical molecules in the CXCL12 – CXCR4 signaling that are involved in the maintenance of self-renewal and metastatic potential of CSCs and miCSCs, a protein – protein interaction (STRING) network analysis of the CXCL12 – CXCR4 axis (green) revealed relevant factors involved in metastasis (red), stemness (purple), sonic hedgehog signaling (blue), AKT signaling (grey) and NF-κB signaling (yellow) (**Fig. 4A**). This interaction analysis revealed BMI1 (polycomb group RING finger protein 4 or RING finger protein 51) as a central player that funnels signals through the CXCL12 – CXCR4 axis, which could provide an explanation for the relationship between EMT and stemness in cancer cells.

**Figure 4.**
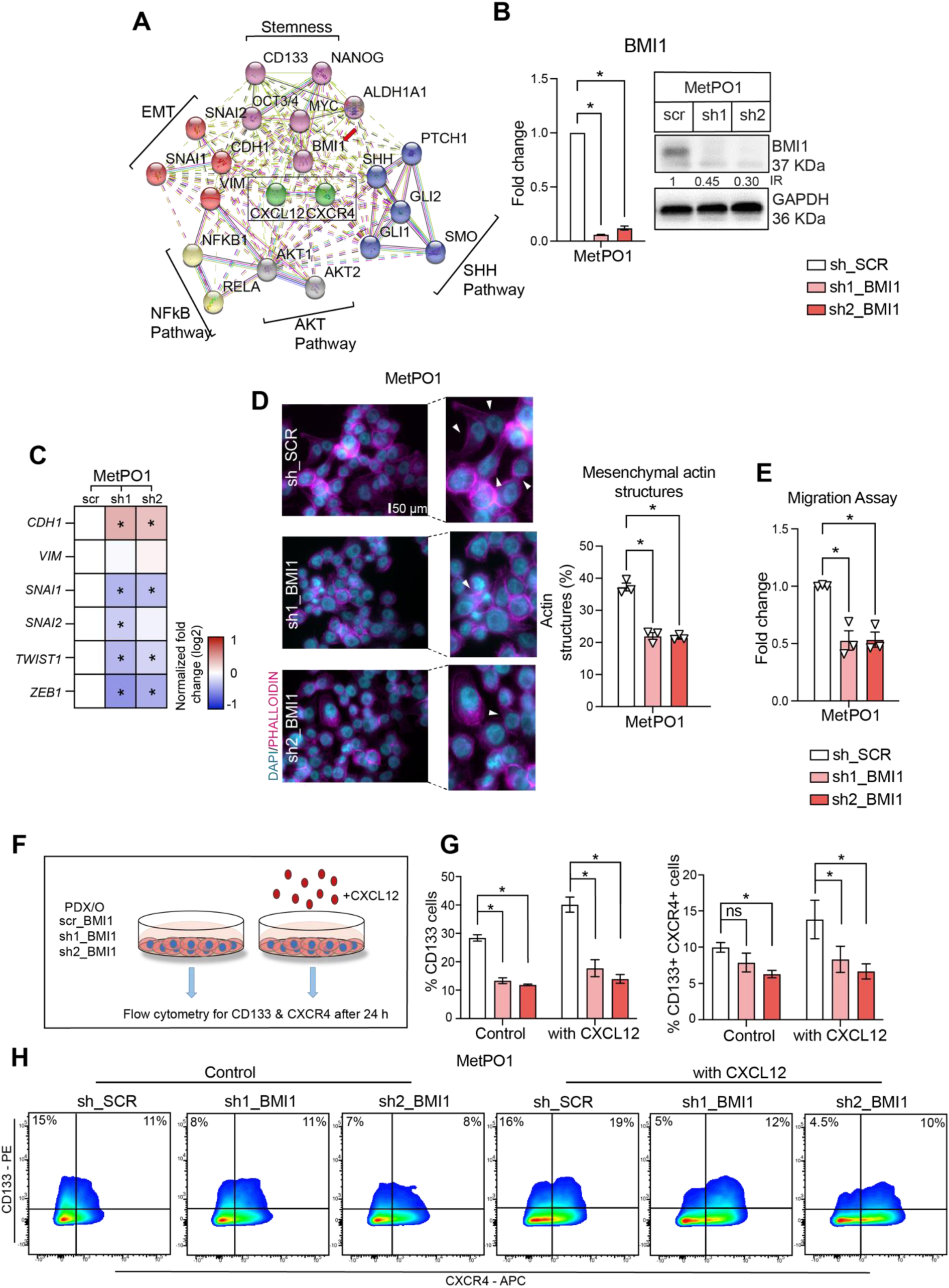
BMI1 downstream CXCL12 – CXCR4 regulates EMT and stemness. (A) Protein – protein interactions of CXCL12 and CXCR4 with relevant factors involved in metastasis (red), stemness (purple), sonic hedgehog signaling (blue), AKT signaling (grey) and NFκB pathway (yellow) (STRING). (B) *BMI1* gene expression analysis and western blot analysis. GAPDH was used as a loading control. (C) Gene expression analysis with genes involved in EMT using qRT-PCR. (D) Immunofluorescence quantifications and representative micrographs for indicated cell lines with white arrowheads marking mesenchymal structures of actin filaments stained with Phalloidin (pink) and nucleus stained with DAPI (blue). (E) Migration assays towards serum containing media. (F) Experimental scheme to evaluate CD133 and CXCR4 surface expression using flow cytometry for Panc354 and MetPO1 (sh_*SCR*, sh1_*BMI1* and sh2_*BMI1*) with (or without) CXCL12. (G) Flow cytometry analysis of CD133+ cells and CD133+ CXCR4+ cells in MetPO1 (sh_*SCR*, sh1_*BMI1* and sh2_*BMI1*) treated with (or without) CXCL12. (H) Representative cytometry plots for MetPO1 cell line. Error bars represent the standard deviation. n≥3 for all experiments. *p < 0.05, ns = not significant.

BMI1 is part of the PRC1 (Polycomb repressive complex) complex and, together with the PRC2 component EZH2, is involved in the maintenance of self-renewal and stemness. In order to delineate the role of BMI1 in CXCL12 – CXCR4-mediated EMT, shRNA-mediated knockdown of BMI1 was performed (**Fig. 4B**; **Supp. Fig 2A**), resulting in an induction of a more epithelial phenotype, as shown by increased CDH1 expression and suppression of crucial EMT markers such as *SNAI1*, *SNAI2*, *TWIST1* and *ZEB1* (**Fig. 4C**; **Supp. Fig. 2E**). Next, phalloidin staining to monitor changes in actin filament architecture (**Fig. 4D**). Interestingly, the actin stress fibers and mesenchymal structures (white arrows) in metastatic MetPO1 sh_SCR cells were drastically impaired upon BMI1 knockdown, resulting in a more epithelial phenotype. Statistically, BMI1 knockdown showed a significant decrease in mesenchymal actin structures in MetPO1 cells upon loss of BMI1. These findings were further functionally substantiated by migration assays, which revealed a significantly decreased migratory capacity of cancer cells after BMI1 knockdown (**Fig. 4E**; **Supp. Fig 2F**).

Having demonstrated the impact of BMI1 abrogation on EMT, we sought to delineate the contribution of BMI1 on the CXCL12 - CXCR4-mediated maintenance of CSCs and miCSCs. Therefore, PDX/PDO BMI1-KD cells were treated with vehicle or CXCL12 for 24 hours and then analyzed for CD133 and CXCR4 expression (**Fig. 4F**). In line with our hypothesis, we found a significant decrease in the CD133+ CSC population after BMI1 knockdown **(Fig. 4G-H**; **Supp. 2H**). Similarly, a significant decrease in the miCSC (CD133+ CXCR4+) population was also observed in BMI1-KD MetPO1cells (**Fig. 4G-H**; **Supp. 2H**). This effect could not be documented in non-metastatic Panc354 cells after BMI1 knock-down (**Supp. 2H**), arguing for a more important role of BMI1 in metastatic cells. Altogether, these results not only demonstrate the significance of CXCL12 – CXCR4 signaling in the maintenance of EMT and stemness features, but also the indispensability of BMI1 in the regulation and maintenance of CSC and miCSC populations.

### JM#21, a potent EPIX4 derivative to target miCSCs

The CXCL12 – CXCR4 axis regulates a bidirectional tumor-stromal signaling loop that ultimately promotes CSCs and miCSCs and their metastatic and chemoresistant phenotypes. Therefore, targeting CXCR4 may provide a significant improvement in the anticancer therapeutic efficacy. The **E**ndogenous **P**eptide **I**nhibitor of C**X**CR**4** (EPI-X4) is a natural antagonist of CXCR4, previously discovered in a human peptide library screen (Zirafi et al., 2015). EPI-X4 derivatives with increased CXCR4-antagonizing activity have been developed and shown to prevent atopic dermatitis and airway inflammation in mouse models (Zirafi et al., 2015). However, analyses showed that JM#21 is not stable in serum-containing culture conditions (**Supp. Fig. 2K**). Therefore, the next experiments were performed using B27 as a serum substitute. In order to uncover the potential of EPI-X4 and its optimized derivatives to functionally target CXCR4 in human PDAC cells (i.e. CXCR4-mediated migratory and metastatic activity), we used 3D transwell migration assays with CXCL12 as chemoattractant to test the inhibitory capacity of EPI-X4 and its modified derivates: WSCO2 (Zirafi et al., 2015), JM#21 (Harms et al., 2021) as well as an inactive variant (Zirafi et al., 2016) included as a control (**Fig. 5A-B**). Incubation with EPI-X4, WSCO2 or JM#21 significantly suppressed the CXCL12-induced migration of Panc354 up to approx. 50% in a dose-dependent manner. Interestingly, JM#21 was effective at even lower concentrations. As expected, the inactive peptide showed no effect on migration. As JM#21 was the most potent inhibitory peptide, we evaluated its anti-migratory capacity in other PDX-derived primary human PDAC cell lines, namely MetPO1, Bo80 and Gö13 (**Fig. 5C**; **Supp. Fig. 2J**).

**Figure 5.**
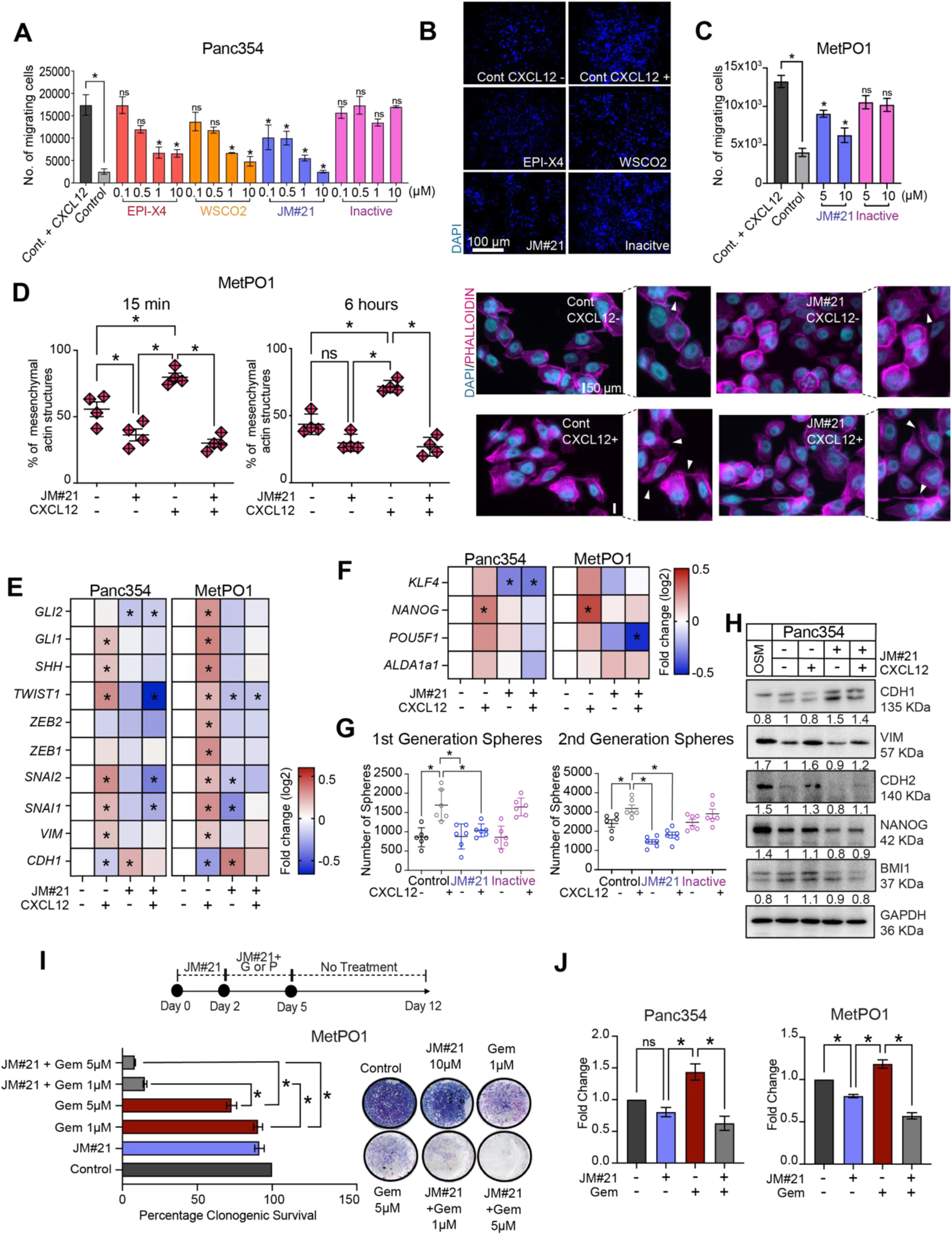
JM#21, most potent EPIX4 derivative to target miCSCs. (A) Migration assays towards CXCL12 using Panc354 for EPI-X4 and its derivatives at depicted concentrations. Pre-treatment with EPI-X4, WSCO2, JM#21 and the inactive peptide was applied for 30 minutes. (B) Representative micrographs (10x, DAPI staining) of transwell migration assays in Panc354 cells for the indicated conditions and concentrations. (C) Migration assays towards CXCL12 for MetPO1 using JM#21 and the inactive peptide at depicted concentrations. (D) Quantifications of percent mesenchymal structures after 15 minutes and 6 hours of CXCL12 treatment in MetPO1 cells. JM#21 pre-treatment was applied for 30 minutes and representative micrographs with white arrowheads marking mesenchymal structures of actin filaments stained with Phalloidin (pink) and nuclear staining using DAPI (blue). (E) Gene expression analysis for indicated cell lines with genes involved in EMT and SHH pathway. (F) Gene expression analysis for indicated cell lines with genes involved in stemness. (G) Sphere formation assays for 1^st^ and 2^nd^ generation of sphere formation. (H) Western blot analysis of CADHERIN-1, VIMENTIN, CADHERIN-2, NANOG and BMI1 for indicated cell lines. GAPDH was used as a loading control. (I) Experimental design for combination therapy analyzing relapse using JM#21, gemcitabine (labelled as G) and paclitaxel (labelled as P). Quantification of cell viability and representative pictures for clonogenic assays after treatment with JM#21 (10*μ*M), gemcitabine (Gem) for indicated concentrations as depicted in experimental design in MetPO1 cell line. (J) Flow cytometry for CD133 in Panc354 and MetPO1cells for the indicated treatments shown as fold change. Error bars represent the standard deviation. n≥3 for all experiments. *p < 0.05, ns = not significant.

We then analyzed whether the inhibition of migration by JM#21 also translated into morphological changes in the cells. The mesenchymal phenotype acquired during EMT is associated with substantial changes in cytoskeletal dynamics and composition. To monitor these changes, we used phalloidin staining of MetPO1 cells subjected to JM#21 treatment and then incubated with CXCL12 (**Fig. 5D**). Incubation with CXCL12 was associated with a swift increase in actin stress fibers formation, mesenchymal structures and adoption of a spindle-like cell shape (white arrowheads), which lasted up to 6 hours (**Fig. 5D**). In line with the migration assay data, JM#21 inhibited the CXCL12-induced formation of mesenchymal structures in MetPO1 cells. In order to characterize the molecular events supporting these findings, we performed pathway-focused gene expression analysis. As expected, the analysis revealed the CXCL12-mediated upregulation of key factors implicated in EMT. Importantly, JM#21 strongly inhibited both baseline and CXCL12-induced expression of several EMT markers such as *SNAI1*, *SNAI2*, *ZEB2*, *TWIST1* (**Fig. 5E**, **Supp. Fig. 3B**). Since previous studies reported the role of sonic hedgehog (SHH) signaling in EMT (Xu et al., 2014; Xu et al., 2012), we analyzed expression levels of SHH pathway-related genes, discovering that JM#21 suppresses expression of *SHH*, *GLI1* and *GLI2* induced by CXCL12 (**Fig. 5E**, **Supp. Fig. 3B**). Furthermore, we found a decrease in CXCL12-induced expression of the stemness factors *POU5F1*, *NANOG*, *KLF4* and *ALDH1a1*, (**Fig. 5F**; **Supp. Fig. 3A**). In line with these findings, both baseline and CXCL12-induced sphere formation of Panc354 cells was inhibited by JM#12 (**Fig. 5G**; **Supp. Fig. 3C**). In summary, these results illustrate the capacity of JM#21 to abrogate CXCL12-induced EMT and stemness in primary PDAC cells.

As expected, the regulation of EMT markers also translated to protein expression: we observed a decrease of CADHERIN1 together with elevated levels of VIMENTIN in response to CXCL12 (**Fig. 5H**; **Supp. Fig. 3D**). Since Oncostatin M (OSM) is a well-established inducer of EMT in pancreatic cancer cells (Alonso-Nocelo et al., 2023; Smigiel et al., 2017), we employed OSM as a positive control for our experiments since CXCL12 showed similar, though less potent effects on EMT. JM#21 treatment resulted in a significant suppression of CXCL12-induced VIMENTIN and CADHERIN-2 expression and induction of CADHERIN-1.

We then evaluated the therapeutic potential of JM#21. Clonogenic assays of Panc354 cells pre-treated with CXCL12 revealed that JM#21 had no effect on cell survival (**Supp. Fig. 3E**). In order to test JM#21 in an in vitro combination therapy of PDAC cells **(Fig. 5I**), PDX/PDO-derived cells cultured with CXCL12 were subjected to 2 days of JM#21 pre-treatment before a 3-day co-treatment with JM#21 and either gemcitabine or paclitaxel. No further treatment was administered for the next 7 days in order to evaluate cell re-growth. Interestingly, single agent treatment with JM#21 or either chemotherapeutic agent did not result in permanent elimination of the treated cells (**Fig. 5I**; **Supp. Fig. 3F-H**). However, combination therapy resulted in significantly impaired colony formation even at lower concentrations of gemcitabine, suggesting a clear benefit of the proposed combination therapy. Since we have previously shown the role of CSCs in chemoresistance and the benefit of combination therapy for PDAC *in vitro* and *in vivo* (Hermann et al., 2007; Hermann et al., 2013; Lonardo et al., 2011; Mueller et al., 2009), we proceeded to evaluate the role of CSCs in the above-mentioned combination treatment and used flow cytometry to quantify CD133+ CSCs after 12 days of treatment (**Fig. 5J**, **Supp**. Fig. 3I). Our analysis revealed that gemcitabine treatment alone was associated with a significant increase in the CD133+ population in relapsed cells, in line with previous data from our laboratory (Hermann et al., 2007).

Interestingly, JM#21 administration decreased the CD133+ population, albeit less pronounced. The suppression of CD133+ population was boosted by the combination treatment. In summary, our data suggest that JM#21 can sensitize chemoresistant PDAC cells and especially CSCs, and that a combination treatment of JM#21 and chemotherapy could be beneficial for treatment.

### Serum stable JM#21 reduces miCSCs in co-cultures with PSCs

Since JM#21 is not stable in serum containing conditions, we set out to facilitate its activity and increase JM#21’s possible translational potential. To address this, we used mesoporous silica nanoparticles that gradually release the peptide, protecting it from proteolytic degradation, to further assess the therapeutic potential of this new CXCR4 antagonist JM#21. JM#21 was encapsulated in mesoporous silica nanoparticles with a radial (MSN) or dendritic (DMSN) oriented pore system (Beitzinger et al., 2021; Rosenholm et al., 2009) (**Fig. 6A**). The effect of nanoparticle encapsulated JM# on cell migration was tested, as described above, and indeed, silica nanoparticle (Si-NP)-encapsulated JM#21 (MSN_JM#21 and DMSN_JM#21) showed substantial suppression of the migratory capacity of all cell lines towards CXCL12 in comparison to empty silica nanoparticles (MSN_Empty and DMSN_Empty). In contrast, and as expected due to the presence of serum, JM#21 alone failed to influence the migration capacity due to the presence of serum (**Fig. 6B-C**; **Supp. Fig. 2K**). The robust decrease in migration confirmed the stability of Si-NP encapsulated JM#21 in serum conditions.

**Figure 6.**
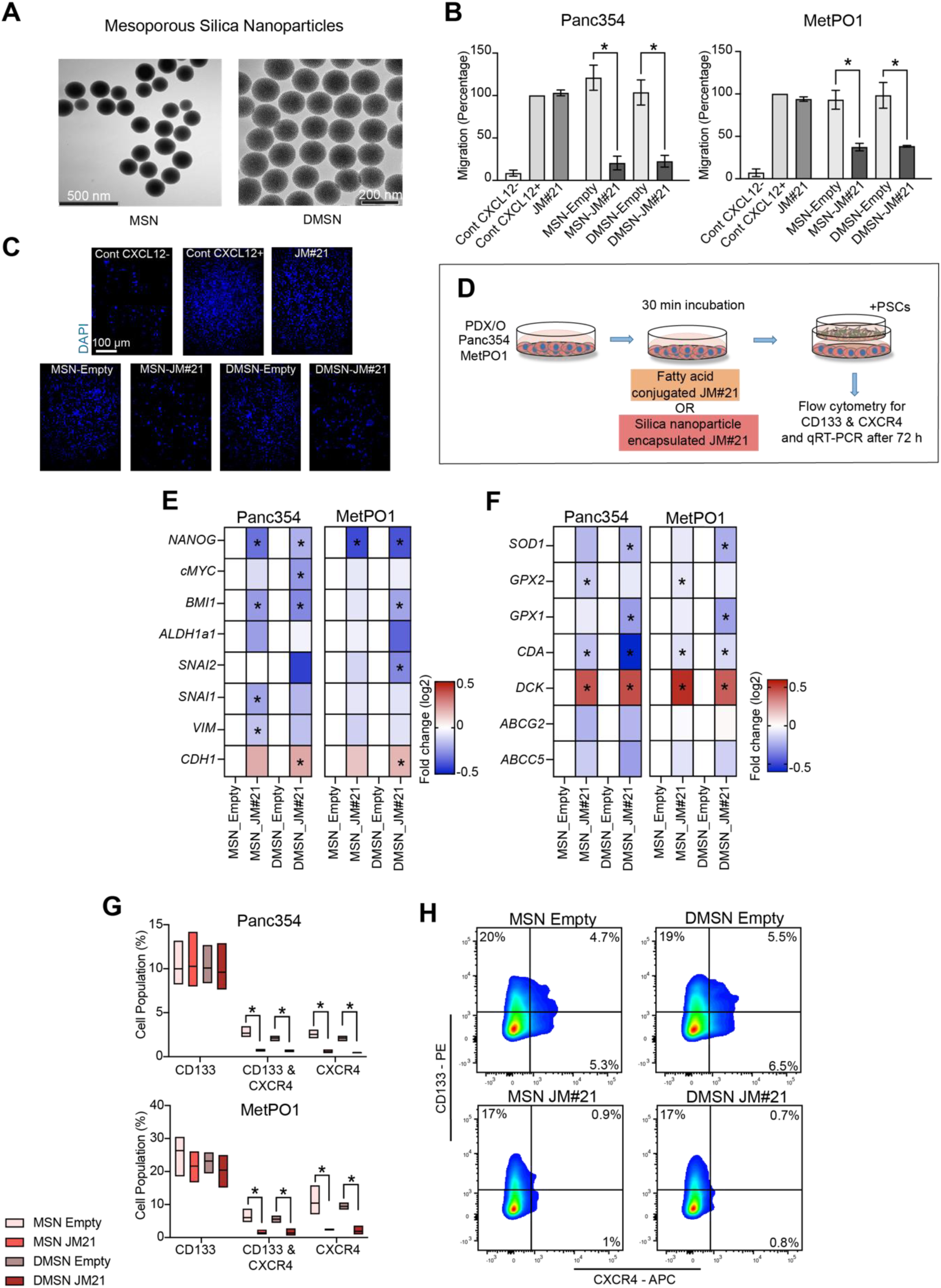
Serum stable JM#21 reduces miCSCs in co-cultures with PSCs. (A) TEM micrographs of silica nanoparticles (MSN and DMSN). (B) Migration assays towards CXCL12 in FBS-containing medium with the indicated compounds tested in Panc354 and MetPO1. (C) Representative micrographs (10x, DAPI staining) of transwell migration assays in MetPO1 cell line. (E) Gene expression analysis for indicated cell lines with genes involved in EMT and stemness after treatment with DMSN and MSN (Empty & JM#21). (F) Gene expression analysis for indicated cell lines with genes involved in chemoresistance and antioxidant signaling after treatment with DMSN and MSN (Empty & JM#21). (G) Flow cytometry analysis performed for CD133+, CD133+CXCR4+ and CXCR4+ cells for depicted cell lines after treatment with DMSN and MSN (Empty & JM#21). (H) Representative cytometry plots for MetPO1 cells after treatment with DMSN and MSN (Empty & JM#21) for CD133+, CD133+CXCR4+ and CXCR4+ cells. n≥3 for all experiments. *p < 0.05, ns = not significant.

We next studied the effect of Si-NP encapsulated-JM#21 pre-treatment of PDX/PDO cell lines co-cultured with PSCs (**Fig. 6D**). The treatment effects were assessed via pathway-focused gene expression analysis involving key genes involved in EMT and stemness (**Fig. 6E**). Treatment with Si-NP-encapsulated JM#21 resulted in a substantial decrease in *NANOG* and *BMI1* gene expression. Furthermore, DMSN_JM#21 significantly increased *CDH1* expression, concomitant with a reduction of *SNAI2* levels, indicating the induction of an epithelial phenotype. In addition, a significant decrease in *CDA1* and increase in *DCK1* expression was observed for both cell lines treated with Si-NP-encapsulated JM#21 compared to empty Si-NP (**Fig. 6F**). Furthermore, a decrease in gene expression of antioxidant enzymes *SOD1*, *GPX1*, *GPX2* was also observed (**Fig. 6F**). Finally, although Si-NP encapsulated JM#21 showed no significant changes in CD133+ expression in flow cytometry analysis, CD133+ CXCR4+ double expression and CXCR4+ expression was substantially reduced (**Fig. 6G-H**). In summary, we were able to show that serum stable JM#21 abrogated the miCSC population in PDX/PDO – PSCs co-culture, and may well provide a novel therapeutic approach to inhibit metastasis in PDAC.

## Discussion

miCSCs play a pivotal role in malignant tumors as they have been reported to lead the metastatic process in the invasive front and are exclusively responsible for metastasis (Cespedes et al., 2018; Hermann et al., 2007; Marechal et al., 2009; Tu et al., 2017; Zhang et al., 2012). We previously identified in PDAC this migrating CSC population as expressing CD133 and CXCR4 (Hermann et al., 2007). Furthermore, CD133+CXCR4+ cells are associated with poor prognosis in patients with colorectal cancer and non-small cell lung carcinomas due to the activation of EMT programs. However, the exact mechanisms that regulate miCSCs are still poorly understood.

Here, we demonstrate that CXCL12 – CXCR4 signaling is a key axis in maintaining miCSC properties. Our findings indicate that elevated secretion of CXCL12 within PSC – PDX/PDO co-cultures confers both mesenchymal and stem-like features to the tumor cell compartment. In the context of tumor microenvironment, the stroma consisting of high number of pancreatic stellate cells are contributing to maintenance of CSCs and miCSCs state. Although PSCs secrete a plethora of cytokines (e.g., TGFß, IL6, Nodal etc.) that promote EMT and stem cell-like properties (Erkan et al., 2012; Lonardo et al., 2012), our data show that abrogation of CXCR4 in PDX/PDO cells is sufficient to disrupt the CXCL12 – CXCR4 axis, despite the presence of recombinant CXCL12 or in co-culture with PSCs. This ultimately translates into the abrogation of mesenchymal features, stem-like properties and chemoresistance, suggesting that CXCL12 released within the TME indeed plays an indispensable role in the maintenance of CSCs and miCSCs.

Mechanistically we show that CXCR4 knockdown not only affects CXCL12-induced EMT initiation, but also the chemoresistance capacity of PDX/PDO cells and CSCs. The CXCL12 – CXCR4 axis interferes with crucial signaling pathways such as PI3K and NF-kB via phosphorylation of AKT and IκB-α, respectively. Upon activation of these pathways, CXCL12 – CXCR4 signaling participates in many biological and physiological functions such as cell migration, proliferation, angiogenesis, release of antioxidant enzymes and chemoresistance (Kato et al., 2022; Seemann and Lupp, 2015; Shen et al., 2013; Zhang et al., 2018). Importantly, our protein-protein interaction analysis revealed BMI1 (Molofsky et al., 2003), as a key component downstream of CXCL12 – CXCR4 signaling, involved in the maintenance of mesenchymal and stem-like features. BMI1 is a member of the polycomb-repressive complex 1 (PRC1) and plays a crucial role in self-renewal through repression of the *INK4A*– *ARF* locus (Valk-Lingbeek et al., 2004; Widschwendter et al., 2007). Abrogation of BMI1 in the patient organoid-derived metastatic primary cell line MetPO1 was associated with decreased phalloidin immunoreactivity of the actin filaments and spindle-like structures. In addition, BMI1 knockdown impaired CSCs and miCSCs, an effect that could not be rescued upon addition of recombinant CXCL12.

Intriguingly, while BMI1 deletion correlated with suppression of CD133 expression, it did not affect sphere formation. At the same time, the ability of BMI1 to control self-renewal and govern key metabolism-associated processes (gluconeogenesis, TCA cycle), particularly in the CD133+ population, was previously reported in glioblastoma multiforme (Vora et al., 2019). Furthermore, BMI1 has also been shown to transcriptionally regulate SOX2 genes in cervical cancer (Mirzaei et al., 2022). These findings suggest that BMI1 function is governed by a restricted number of stemness-associated factors reasoning the incapacity of BMI1 knock-down to impact sphere formation in our experimental setting. Additionally, the role of other compensatory pathways for the loss of BMI1 at the level of self-renewal cannot be ruled out. Importantly, BMI1 has been shown to potentiate self-renewal capacity through the activation of PI3K/AKT and SHH/GLI1 signaling pathways either in a direct manner by induction of Nanog/NF-κB, or indirectly upon hyperactivation of the PI3K/Akt/NF-κΒ axis (Jiang et al., 2012; Liu et al., 2006; Song et al., 2009). We have previously described that the MEK/ERK axis promotes migratory features of CSCs in PDAC (Walter et al., 2019). In line with these findings, we were able to demonstrate here that BMI1 deletion impaired ERK1/2 phosphorylation, further underscoring the role of BMI1 in (mi)CSC maintenance.

From a therapeutic perspective, Yin and colleagues showed that BMI1 inhibition is associated with a sensitization of PDAC cells towards gemcitabine (Yin et al., 2016). Interestingly, our PDX/PDO-derived cells in PSC co-culture or challenged with recombinant CXCL12 displayed a sustained BMI1 activation. At the same time, CXCR4 suppression led to pronounced reduction in BMI1 protein expression, which could not be restored by CXCL12 or only partially by PSC co-culture. These findings establish BMI1 as a funnel for CXCL12- mediated effects on stemness and migration. Altogether, our study demonstrates that the CXCR4/BMI1 axis is a potential target for future therapeutic interventions aiming at suppression of CSCs and miCSCs. As such, we employed endogenous human peptides as a novel therapeutic strategy to target miCSCs-mediated migration and metastasis. This study reveals for the first time the capability of EPI-X4 and its potent derivative JM#21, to efficiently block CXCR4 with a subsequent suppression of migrating CSCs in PDAC. Furthermore, JM#21 appears to be not only a highly specific and remarkably active CXCR4 antagonist, but also a compound capable of substantial inhibition of CXCL12-driven tumor cell migration and sphere formation, opening a novel avenue to improve PDAC treatment.

Gemcitabine treatment predominantly affects differentiated cells while having virtually no effect on CSCs. Both in previous studies and in the current work, we have shown that gemcitabine treatment is associated with expansion of drug-resistant CD133+ cancer stem cells but also of CD133+CXCR4+ miCSCs (Cash et al., 2020; Hermann et al., 2007; Hermann et al., 2013; Mueller et al., 2009). The effectiveness of combination therapies targeting specific stemness-associated pathways together with chemotherapy was previously documented (Gallmeier et al., 2011; Hermann et al., 2013; Lonardo et al., 2011; Mueller et al., 2009; Sainz et al., 2015). Strikingly, a combination therapy of gemcitabine or paclitaxel together with the novel CXCR4-antagonist JM#21 re-sensitized drug-resistant PDAC cells to chemotherapy. Furthermore, the use of serum-stable, nanoparticle-encapsulated JM#21 in co-culture settings impaired the expression of genes that regulate chemoresistance (e.g., *CDA1, ABCG2, ABCC5*) as well as BMI1. These findings further extend the knowledge on the ability of JM#21 to reduce CSC burden in addition to blocking CXCR4-mediated migration and making chemoresistant cancer cells vulnerable to conventional therapies. In summary, we show herein for the first time that serum-stable endogenous peptides such as JM#21 represent a potent novel therapeutic entity, which robustly targets CXCR4 to reduce migration and eliminate miCSCs in PDAC.

## Materials and Methods

### Primary cell lines and co-culture

All cell lines were maintained at 37°C in a humidified atmosphere with 5% CO2 in the indicated culture media (DMEM for Panc215, DMEM:F12 for PSCs and RPMI for rest) supplemented with 10% FBS (ThermoFischer Scientific, 10270-106) (20% FBS + 10% PSCs - CM for pancreatic stellate cells), 1% Penicillin-Streptomycin (PAN Biosystems, P06-07100) and 1% Glutamine (Gibco, 41965-039). For experiments, cells were used in early passages and were recovered from frozen stocks on a regular basis. Single cells were counted with Neubauer chambers and seeded accordingly.

Primary human pancreatic xenografts (PDXs) were generated from resected excess pancreatic carcinoma tissue that was subcutaneously implanted into nude mice in vivo. Primary human pancreatic xenograft-derived cell lines (PDX) were derived from excised PDXs as per published protocols (ref). The patient derived organoid cell lines (PDO) MetPO1 and MetPO2 originate from resected excess liver metastasis grown as organoids in vitro. Bo80 cells were kindly provided by S. Hahn. Gö5, Gö7 and Gö13 cell lines were kindly provided by E. Heßmann. Primary PSCs were kindly provided by T. Seufferlein.

Co-culture experiments with PSCs: PDX/PDO derived cell lines were cultured in 6-well plates at a density of 1 x 10^5^ cells per well. PSCs were cultured in inserts with 1μm pore size PET membranes at a density of 1 x 10^5^ cells per insert. Cells were kept separate at 37°C in a humidified atmosphere with 5% CO2 in the respective culture media for 24 hours. Subsequently, media for both 6-well plate and inserts were replaced with RPMI (with 10% FBS, 1% Pen-Strep and 1% Glutamine) and inserts with PSCs were placed over PDX/PDO derived cell lines in 6-well plates and incubated additionally for 72 hours to establish dual cultures. Control PDX/PDO-derived cells were cultured at the density of 1 x 10^5^ cells per insert.

### Conditioned media for PSCs maintenance

PSCs were thawed and cultured in a T75 flask in DMEM:F12 (1:1) together with 20% FBS, 1% Pen-Strep and 1% Glutamine for 7 days at 37°C in a humidified atmosphere with 5% CO2. Media was collected in a 50ml tube and centrifuged at 1000 rpm for 5 minutes to remove debris. Media was next filter-sterilized (0.45μm filter), labelled and stored as PSCs conditioned media (PSCs - CM) at 4°C.

### Primed and non-primed PSCs – Conditioned Media (CM) preparation

PDX/PDO-derived cell lines and PSCs were cultured separately in 10cm dishes with their respective media for 24 hours at 37°C in a humidified atmosphere with 5% CO2. Subsequently, media from PDX/PDO-derived cell lines was collected in a 50ml tube and centrifuged at 1000 rpm for 5 minutes to remove debris. Media was next filter-sterilized (0.45μm filter) and was labelled as PDX/PDO – CM. PSCs media was replaced with this PDX/PDO – CM and was allowed to grow for additional 24 hours to prime or activate the PSCs. After 24 hours media from primed PSCs was collected in a 50ml tube and centrifuged at 1000 rpm for 5 minutes to remove debris. Media was next filter-sterilized (0.45μm filter) and was labelled as ‘Primed’ PSCs – CM. ‘Non-primed’ PSCs – CM was prepared by removing the PDX/PDO – CM based priming step and adding fresh RPMI (with 10% FBS, 1% Pen-Strep and 1% Glutamine) media to PSCs culture dish instead.

### Sphere Culture

For first-generation spheres, following treatment or co-culture, 10,000 cells per milliliter were seeded in ultralow attachment plates (Corning, 3473) and cultured at 37°C in a humidified atmosphere with 5% CO2. Spheres were cultured in DMEM-F12 (Thermo Fisher Scientific, 10565018) supplemented with B-27 (Thermo Fisher, 17504044) and basic fibroblast growth factor (Novoprotein, CO46). After 7 days of incubation, > 40μm and < 120μm were quantified using CASY TT (OMNI Life Science, 5651697). For second-generation spheres, after counting the first-generation spheres, they were disrupted using trypsin for 15 minutes at 37°C to make single cells. Equal number of cells per milliliter were seeded back in ultralow attachment plates. After 7 days of incubation, > 40μm and < 120μm were quantified using CASY TT.

### Clonogenic Assay

Cells were seeded at a density if 10^4^/ml in a 24-well plate. After treatment, media was removed, and the wells were washed three times with PBS. After washing, 1 ml of diluted Giemsa solution (Merck, 1.09204.0100) (1: 20 in PBS) was added to each well and incubated for 20 min on a shaker with 150 rpm. Later, Giemsa solution was removed, and the wells were rinsed with water slowly. Dried plates were then used for imaging. For quantification, 10% acetic acid was added to Giemsa-stained cells and incubated for 20 min on a shaker with 150 rpm for blue color to develop. 100*μ*l of this solution was then transferred into a 96- well plate, absorbance was measured using Infinite 200 PRO plate reader (Tecan, Switzerland) at 590 nm.

### Migration Assay

Migration assays were performed using inserts with 8*μ*m pore size PET membranes (Corning, 353097). Serum (10% FBS) or CXCl12 (10nM) was used in the companion plates as chemoattractant. After 24 hours, invaded cells were fixed with 4% PFA and stained with DAPI (Merck, 10236276001). Ten random 10x fields were chosen and photographed (Keyence), and the pictures were quantified using ImageJ.

### Molecular Cloning

Bacterial culture and plasmid isolation: DH5α-T1R- *E. coli* transformed with BMI1, CXCR4 or scramble targeting shRNA plasmids were streaked onto LB agar plates supplemented with 50mg/ml Ampicillin and incubated overnight at 37°C. A single colony was picked from the plate and transferred to a tube containing LB media. Bacterial growth was facilitated by overnight (*<*18 hours) incubation at 37 °C and 220 rpm in Certomat^®^ R. 1 mL of this starting culture was used to inoculate 250mL LB in a conical flask incubated overnight (*<*18 hours) at 37°C and 220 rpm. Bacterial work was conducted near an open flame. For plasmid isolation, PureLink^TM^ HiPure Plasmid Mediprep Kit from Invitrogen was utilized. LB media containing bacterial growth was centrifuged for 10min at 5000 rpm and 4°C, the plasmids were isolated in accordance with the kits protocol. Purified plasmids were resuspended in 100µL TE buffer and proper isolation was confirmed by plasmid restriction.

Lentiviral particle production in Lenti-X cells: Lenti-X (293T) cells were seeded at 5 x 10^6^ cells per 10cm dish, cultured for 24 hours, and transfected with 8*µ*g plasmids of interest (shBMI1; shCXCR4; shSCR), 5.5*µ*g psPAX2 (lentiviral packaging), 2µg pMD2 (lentiviral envelope) and 8.75% PEI transfection reagent (50 mg PEI, 20 mM HEPES and 150mM NaCl; for DNA charge opposition) in serum-free DMEM. After 4 hours at 37 °C, media was replaced with DMEM supplemented with 10% FBS, 1% Pen-strep and 1% Glutamine. Virus was harvested after 2 and 4 days after transfection, pooled and centrifuged at 1500 rpm, room temperature for 2 min to pellet remaining cells. Supernatants were sterile filtered (0.45*µ*m), mixed with Lenti-X Concentrator (3:1) (Clontech, 631231), and incubated at 4°C for 30 min prior to 45 min centrifugation at 1500 rpm at 4°C for virus precipitation. Virus pellets were resuspended in DMEM.

Lentiviral transduction and selection of target cells: Human PDAC cell lines were seeded at a density of 1 x 10^6^ cells per 2ml in 6 well plates. They were then infected with 0 - 25*µ*l virus polybrene (10*µ*g/ml enhances retroviral infection) containing RPMI after 24 hours, as well as 48 hours of culture. Cell culture media was replaced 24 hours after the second infection and the cells were allowed to recover for 48 hours. For drug selection, media was substituted with puromycin (3mg/ml) supplemented RPMI every three days, until non-transduced cells (0*µ*l virus) in control wells died. Consequently, only successfully transduced cells retaining the puromycin resistance gene survived the treatment. Upon confluency, samples of each transduction were collected for qRT-PCR analysis to assess the minimal virus concentration with the strongest downregulation. Cells with the optimal virus concentration were expanded for conducting experiments.

### RNA isolation and real-time PCR

Total RNA was prepared using the RNeasy kit with on-column genomic DNA digestion following the manufacturer’s instructions (Qiagen, 74106). First-strand cDNA was prepared using the QuantiTect Reverse Transcription Kit (Qiagen, 205314). Reactions were performed with the PerfeCTa SYBR Green FastMix PCR Reagent (Qiagen, 204057) using a QuantStudio^TM^ 3 machine (Applied Biosystems) using primers (Table 1). Results were analyzed using the 2^-ddCt^ method relative to *HPRT* or *GAPDH* (for human primary cell lines). Reactions were carried out from at least three independent experiments.

**Table 1:**
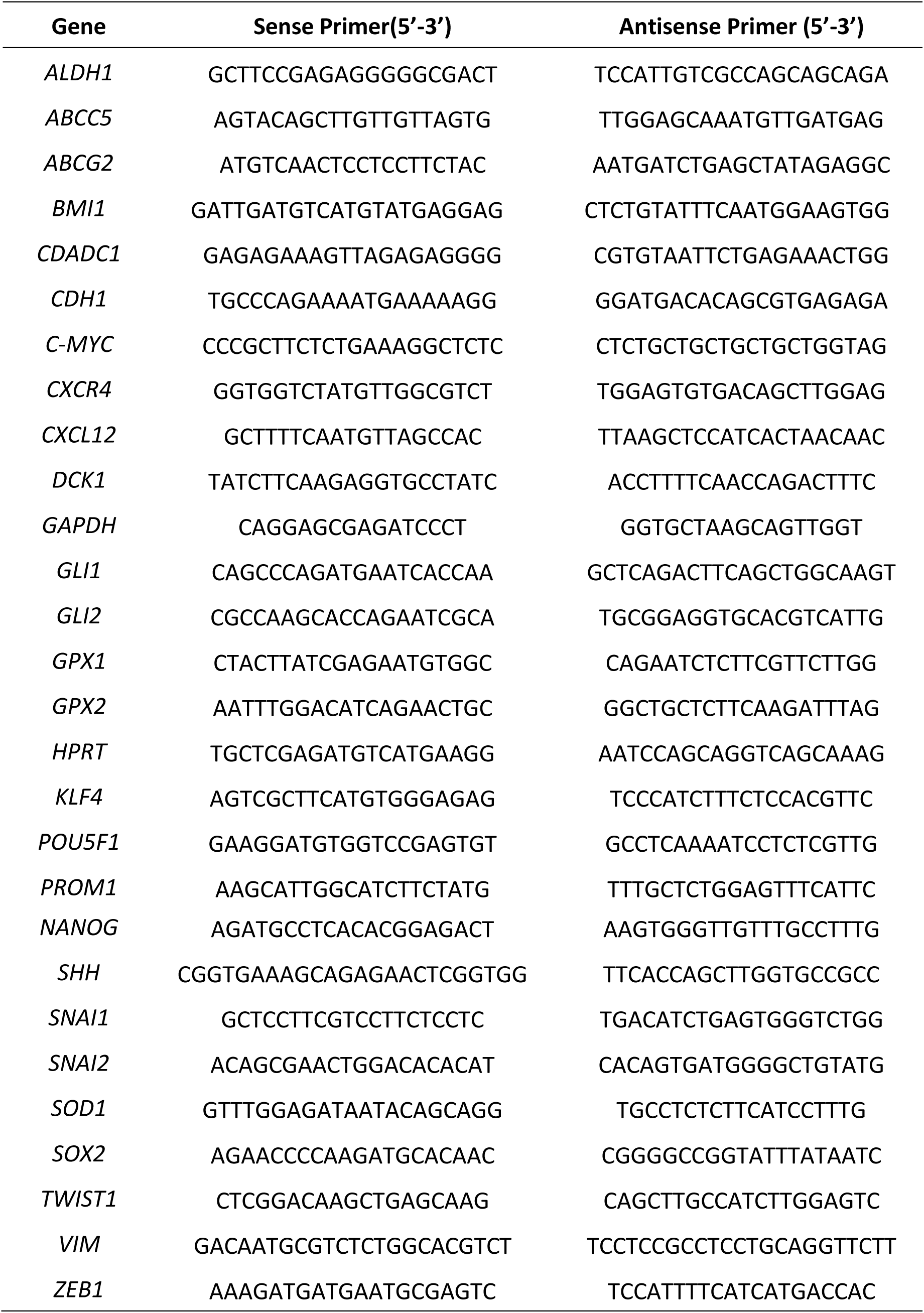
Overview of primers used with their sequences.

**Table 2:** Overview of plasmids used.

| Plasmid | Number | Manufacturer |
| --- | --- | --- |
| pLKO.1-puro-shRNA-BMI1 | TRCN0000020155 | Sigma-Aldrich |
| pLKO.1-puro-shRNA-BMI1(2) | TRCN0000020158 | Sigma-Aldrich |
| pLKO.1-puro-shRNA-CXCR4 | TRCN0000256864 | Sigma-Aldrich |
| pLKO.1-puro-shRNA-CXCR4 (2) | TRCN0000256865 | Sigma-Aldrich |
| pLKO.1-puro-shRNA-scramble | MFCD07785395 | Sigma-Aldrich |
| pMD2.G | 12259 | Addgene |
| psPAX2 | 12260 | Addgene |

### Protein extraction and western blot analysis

Harvested cells were lysed in ice-cold RIPA buffer (Cell Signaling, 9806S) supplemented with PhosSTOp™ (Merck, 4906845001) and a protease inhibitor cocktail (Merck, 11836170001). For each sample, equal amounts of protein were resolved on a 10% SDS-polyacrylamide gel and immunoblotted onto PVDF membranes (GE Healthcare, 10600021). Membranes were blocked for 2 hours in 5% BSA in 1x TBST, probed with the primary antibodies (Table 3) overnight at 4°C, washed with 1x TBST, and incubated with indicated secondary antibody (Table 3) for 2 hours. The chemiluminescence detection was performed according to the manufacturer’s instructions (Merck, WBKLS0500). Stripping of blots was performed when required with Restore^TM^ Western Blot Stripping Buffer according to the manufacturer’s instructions (Thermo Scientific, 21059). Blots were re-blocked after stripping overnight at 4°C in 5% BSA in 1x TBST before probing with the primary antibody.

**Table 3:** Overview of antibodies used for western blot.

| Antibody (Clone) / Species | Dilution | Company | Catalogue# |
| --- | --- | --- | --- |
| **Primary Antibodies** |  |  |  |
| BMI1 (F-9) / Mouse | 1:1000 | Santa Cruz<br>Biotechnology | sc390443 |
| Vimentin (D21H3) / Rabbit | 1:1000 | Cell Signaling | 5741S |
| AKT / Rabbit | 1:1000 | Cell Signaling | 9272 |
| Phosphorylated (Ser473) AKT /<br>Rabbit | 1:1000 | Cell Signaling | 9271 |
| IκB-α / Rabbit | 1:1000 | Cell Signaling | 9242 |
| Phosphorylated (Ser32) (14D4) IκB-α / Rabbit | 1:1000 | Cell Signaling | 2859 |
| Nanog (polyclonal) / Rabbit | 1:1000 | Cell Signaling | 3580S |
| c-MYC (E5Q6W) / Rabbit | 1:1000 | Cell Signaling | 18583S |
| Sox9 (polyclonal) / Rabbit | 1:2000 | Merck | ab5535 |
| N-Cadherin (D4R1H) / Rabbit | 1:1000 | Cell Signaling | 13116S |
| E-Cadherin (24E10) / Rabbit | 1:1000 | Cell Signaling | 3195S |
| Slug (H-140) / Rabbit | 1:1000 | Santa Cruz | sc15391 |
|  |  | Biotechnology |  |
| GAPDH (polyclonal) / Rabbit | 1:10000 | Sigma-Aldrich | G9545 |
| **Secondary Antibodies** |  |  |  |
| Recombinant Anti-mouse IgGκ L | 1:2000 | Santa Cruz | sc516102 |
| (polyclonal), HRP-conjugated |  | Biotechnology |  |
| Goat Anti-rabbit IgG H+L | 1:2000 | Thermo Fisher | G21234 |
| (polyclonal), HRP-conjugated |  | Scientific |  |

### Immunofluorescence

For PSCs: Pancreatic stellate cells (PSCs) were cultured at a density of 50,000 on coverslips in a 6-well plate. PDX/O cell lines were grown in inserts with 1*μ*m pore size PET membranes. After 24 hours, PDX/O cell lines grown in inserts were placed over PSCs on coverslips. After 72 hours, pancreatic stellate cells were washed three times with PBS.

For PDX/PDO cell lines: PDX/PDO cell lines were cultured at a density of 50,000 on coverslips in a 6-well plate. After 24 hours of treatment, cells were washed three times with PBS.

After washing cells were fixed with 2% PFA (Sigma, 158127) for 20 min at room temperature and then washed again three times with PBS. Next PSCs were permeabilized with 0.7% TritonX-100 (Fluka, 93420) solution for 15 min at room temperature followed by three PBS washing. Coverslips with PSCs were then incubated with Nile red (1:500 diluted in PBS) (Sigma, N3013) and Phalloidin-Atto (1:500 diluted in PBS) (Sigma, 94072) inside a dark humid chamber for 60 min. PSCs were washed three times in PBS and mounted on slides using Prolong™ Gold reagent with DAPI. PSCs were visualized with fluorescence at 465nm, 488nm and 565 nm was visualized and photographed EVOS FL (Invitrogen, AMF4300) and PDX/PDO cells visualized with fluorescence at 565 nm was visualized and photographed Zeiss Axio Vert.A1(Zen Blue).

### Flow Cytometry

For surface protein staining, 1 x 10^6^ cells were resuspended in 100µl ice-cold FACS buffer supplemented with 6 µl Gamunex for 15 min at 4°C. Cells were then incubated with CD133 (PE, Miltenyi Biotec, #130-113-108) and CXCR4 (APC, eBioscience, #17-9999-42) antibodies or an appropriate isotype control for 30 mins at 4°C. After a PBS wash, cells were resuspended in 500µl of ice-cold FACS buffer with DAPI in FACS tubes for analysis.

### Enzyme-linked immunosorbent assay (ELISA)

Media levels of secreted CXCL12 were determined using CXCL12A Human ELISA kit (Thermo Fischer Scientific, EHCXCL12A) according to the manufacturer’s guidelines. Conditioned media from PSCs monoculture, PDX/O – PDX/O and PDX/O – PSCs co-cultures were collected after 72 hours and stored at -80°C. Thawed conditioned media samples were diluted 2-fold with assay diluent B. Absorbance was measured using Infinite 200 PRO plate reader (Tecan, Switzerland) at 450 nm.

### Antibody-competition assay

50,000 SupT1 cells/well were seeded in 96-well V-bottom microtiter plates in FACS buffer. Buffer was removed by centrifugation and cells precooled at 4°C for 15 min. In the meantime, compounds were serially diluted in ice cold PBS and 12G5-APC antibody was diluted in cold FACS buffer at a concentration of 0.49nM. Afterwards, 15*µ*l of compound directly followed by 15*µ*l of antibody was added to the cells. For stability experiments, the mixture was carefully mixed after addition to the cells. Cells were incubated at 4°C for 2 hours before unbound antibody and compounds were removed by 2 washing steps followed by fixation in 2% PFA buffer. Cells were analyzed for MFI values of bound antibody by flow cytometry using FACS CytoFLEX. For calculation, the isotype control (Table 4) was subtracted, and values normalized to 12G5 stained PBS control.

**Table 4:**
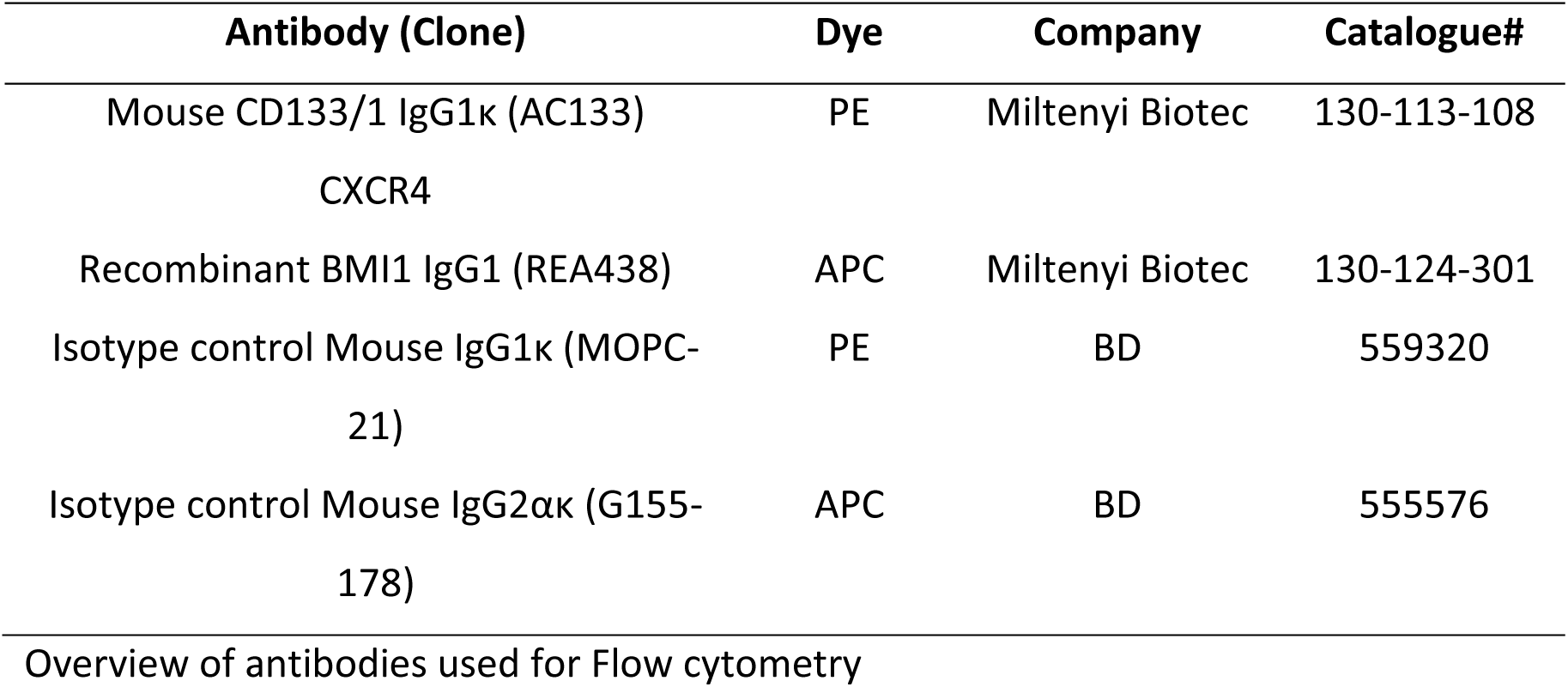

| **Antibody (Clone)** | **Dye** | **Company** | **Catalogue#** |
| --- | --- | --- | --- |
| Mouse CD133/1 IgG1κ (AC133) | PE | Miltenyi Biotec | 130-113-108 |
| CXCR4 |  |  |  |
| Recombinant BMI1 IgG1 (REA438) | APC | Miltenyi Biotec | 130-124-301 |
| Isotype control Mouse IgG1κ (MOPC- | PE | BD | 559320 |
| 21) |  |  |  |
| Isotype control Mouse IgG2ακ (G155- | APC | BD | 555576 |
| 178) |  |  |  |

Overview of antibodies used for Flow cytometry

### Microtiter based stability determination

JM#21 peptide was 150-fold diluted in media supplemented with PBS, 10% FCS or to reach a final concentration of 20*µ*M. The t = 0 sample was immediately taken and stored at -80°C. Media/peptide mixture was then transferred to 37°C and shaken at 350 rpm. At given time points, samples were taken and stored at -80°C. For measuring the functional activity of the media/peptide samples, the mixtures were thawed and serially diluted in ice cold PBS (starting with 100 % sample). 12G5-APC antibody competition was then performed as described before.

### Fatty acid conjugation with JM#21

JM#21 conjugation to palmitic acid (JM#143 and JM#194) has been previously described in the published thesis Harms M: Endogenous CXCR4 Antagonists: Role in HIV-1 Transmission and Therapy of CXCR4-linked diseases (http://dx.doi.org/10.18725/OPARU-41859), (Harms et al., 2021) PhD Dissertation, Institute of Molecular Virology, University of Ulm, 2021.

### Loading JM#21 to silica nanoparticles

The mesoporous silica nanoparticles with a radially (MSN) and dendritic (DMSN) oriented pore system were synthesized as described (Beitzinger et al., 2021; Beitzinger et al., 2024; Rosenholm et al., 2009). In brief, cetyltrimethylammonium bromide (CTAB) was used as a structure directing agent and a mixture of tetramethyl orthosilicate (TMOS) and (3- aminopropyl) trimethoxysilane (APTMS) acted as silica precursors. The final synthesis mixture had a molar ratio of 0.68 TMOS: 0.1 APTMS: 1.00 CTAB: 0.21 NaOH: 1746 MeOH: 4142 H2O. Instead of extraction with acidic ethanol, the surfactant was removed via calcination at 550°C for 5.5 hours. JM#21 was adsorbed onto the radial mesoporous silica nanoparticles (MSN) and dendritic mesoporous silica nanoparticles (DMSN) similar to as described previously in (Beitzinger et al., 2021).

### Bioinformatics prediction

GEPIA (http://gepia.cancer-pku.cn/intex.html) was applied to conduct tumor/normal differential expression analysis (Tang et al., 2017). The protein – protein interactions were carried out using STRING database (Jensen et al., 2009).

### Statistics

Results for continuous variables are presented as mean ± SEM unless stated otherwise or as median with quartiles and min/max values for box plots. Statistical analysis was performed using GraphPad Prism version 9. In order to detect normal distribution, Shapiro-wilk tests were performed. Normally distributed data was analyzed using two-way ANOVA (or one-way ANOVA for knockdown experiments), otherwise Mann-Whitney U test was applied. Fold change data was analyzed using student’s t test. *p values < 0.05 were considered statistically significant. For each experiment, the sample size is indicated in the figure legend.

## Acknowledgements

We gratefully acknowledge Pierre-Olivier Frappart for the support during the generation of the organoid cultures, and we are indebted to Andrea Wissmann for excellent technical support.

## Author contributions

K.T. and P.C.H. conceived the study, designed the experiments, analyzed the data, and wrote the manuscript. A.L., S.I., S.H., M.H., F.A., S.A. and K.W. performed experiments and/or analyzed data. M.H. synthesized the JM#21 peptide. B.B. and R.S. synthesized the JM#21 loaded and unloaded silica nanoparticle formulations. A.K., N.A., T.S., S.H., E.H., B.S.Jr, J.M. and M.L. contributed to the study design, edited and revised the manuscript.

## The Paper Explained

### Problem

Pancreatic ductal adenocarcinoma (PDAC) is one of the most aggressive and lethal cancers, with high metastatic potential and resistance to conventional therapies. A critical aspect driving metastasis in PDAC is the presence of migrating cancer stem cells (miCSCs), characterized by CD133+CXCR4+ expression. These cells play a key role in metastasis and therapeutic resistance, yet effective druggable targets remain largely unknown. Pancreatic stellate cells within the tumor microenvironment secretes CXCL12, which activates CXCR4 on (mi)CSCs, enhancing their stemness, migration and chemoresistance. In this study, we aim to delineate and disrupt this signaling pathway to improve PDAC treatment strategies.

### Results

In this study we identify the transcription factor BMI1 as a crucial downstream effector of CXCL12/CXCR4 signaling. Knockdown of CXCR4 and BMI1 significantly reduces miCSC migration, sphere formation, and EMT, confirming BMI1’s role in sustaining PDAC aggressiveness. We show that the novel CXCR4 inhibitors EPI-X4 and its optimized derivative JM#21 effectively blocks CXCL12/CXCR4 signaling, leading to suppression of EMT, stemness, and self-renewal in PDAC cell lines. Moreover, JM#21 enhances the efficacy of standard chemotherapies (gemcitabine and paclitaxel), making resistant PDAC cells more responsive to treatment.

### Impact

Targeting the CXCL12/CXCR4/BMI1 axis presents a promising therapeutic strategy for PDAC. By inhibiting miCSC activity, JM#21 disrupts key mechanisms of migration and chemoresistance, which could potentially improve patient outcomes. Notably, JM#21 is a novel, optimized derivative of the endogenous peptide EPI-X4, specifically designed to enhance CXCR4 inhibition. This elevated potency and stability make it a viable therapeutic candidate. Integrating peptide therapeutics such as JM#21 into combination therapies, provides an augmented approach for existing treatments and offers an effective and innovative strategy to combat PDAC’s deadly progression.

---

**Supplementary Figure 1.**
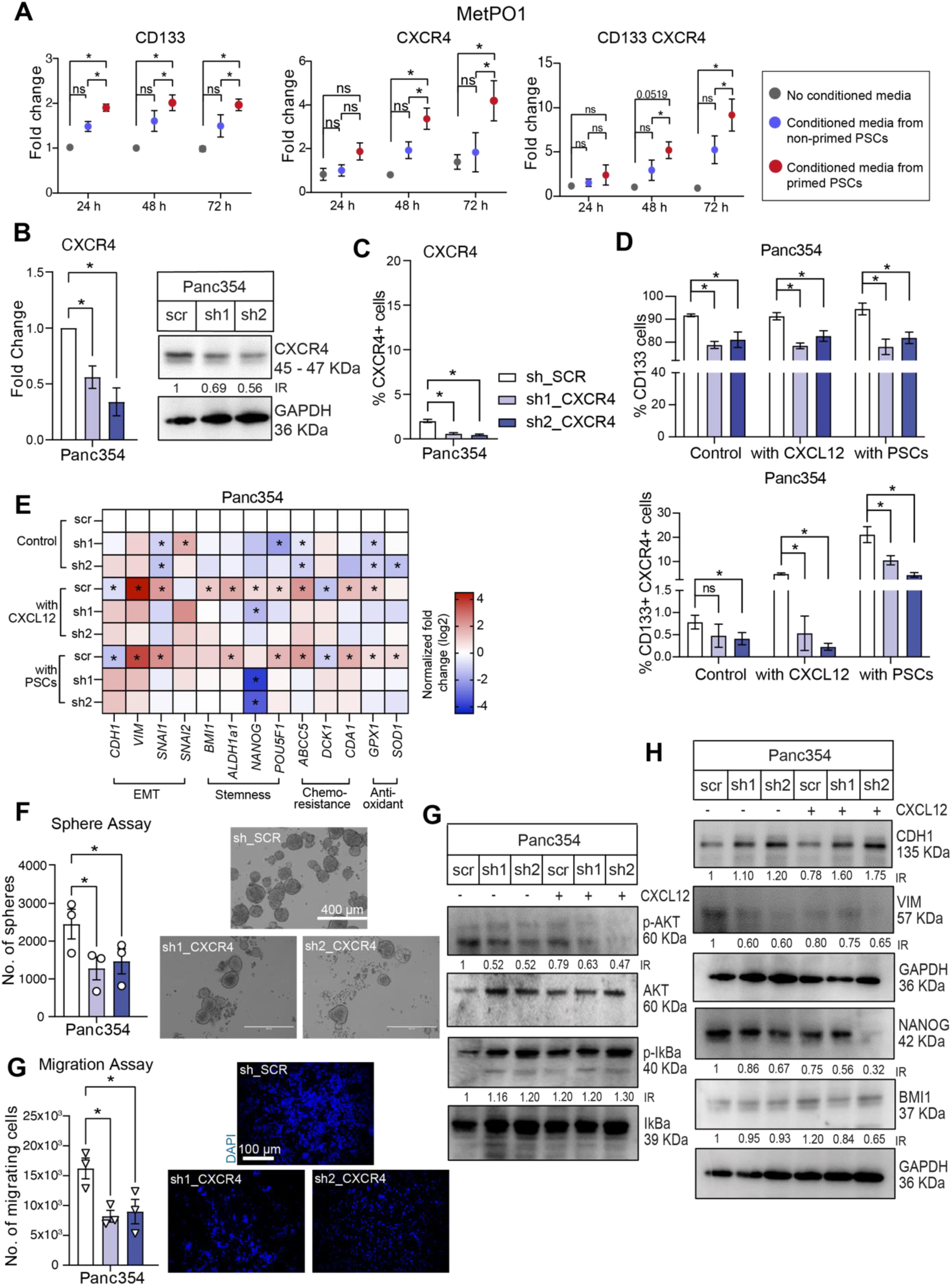
(A) Flow cytometry analysis performed on MetPO1 when exposed to no conditioned media (grey), conditioned media from non - primed PSCs (blue) or conditioned media from primed PSCs (red) for CD133+ cells, CXCR4+ cells and CD133+ CXCR4+ cells represented as fold change against no conditioned media. (B) *CXCR4* gene expression analysis and western blot analysis in Panc354. GAPDH was used as a loading control. (C) Flow cytometry analysis of CXCR4 surface expression. (D) Flow cytometry analysis of CD133+ CSCs and CD133+ CXCR4+ miCSCs in Panc354 (sh_*SCR*, sh1_*CXCR4* and sh2_*CXCR4*) treated with (or without) CXCL12 or co-cultured with PSCs. (E) Targeted gene expression analysis for indicated genes in Panc354 (sh_*SCR*, sh1_*CXCR4* and sh2_*CXCR4*) treated with (or without) CXCL12 or co-culture with PSCs. (F) Sphere formation assay and representative pictures after CXCL12 treatment in Panc354. (G) Migration assay towards CXCL12 and representative pictures in Panc354. (H) Western blot analysis of labelled protein markers for Panc354 cell line and treatment conditions. GAPDH was used as a loading control. Error bars represent the standard deviation. n≥3 for all experiments. *p < 0.05, ns = not significant.

**Supplementary Figure 2.**
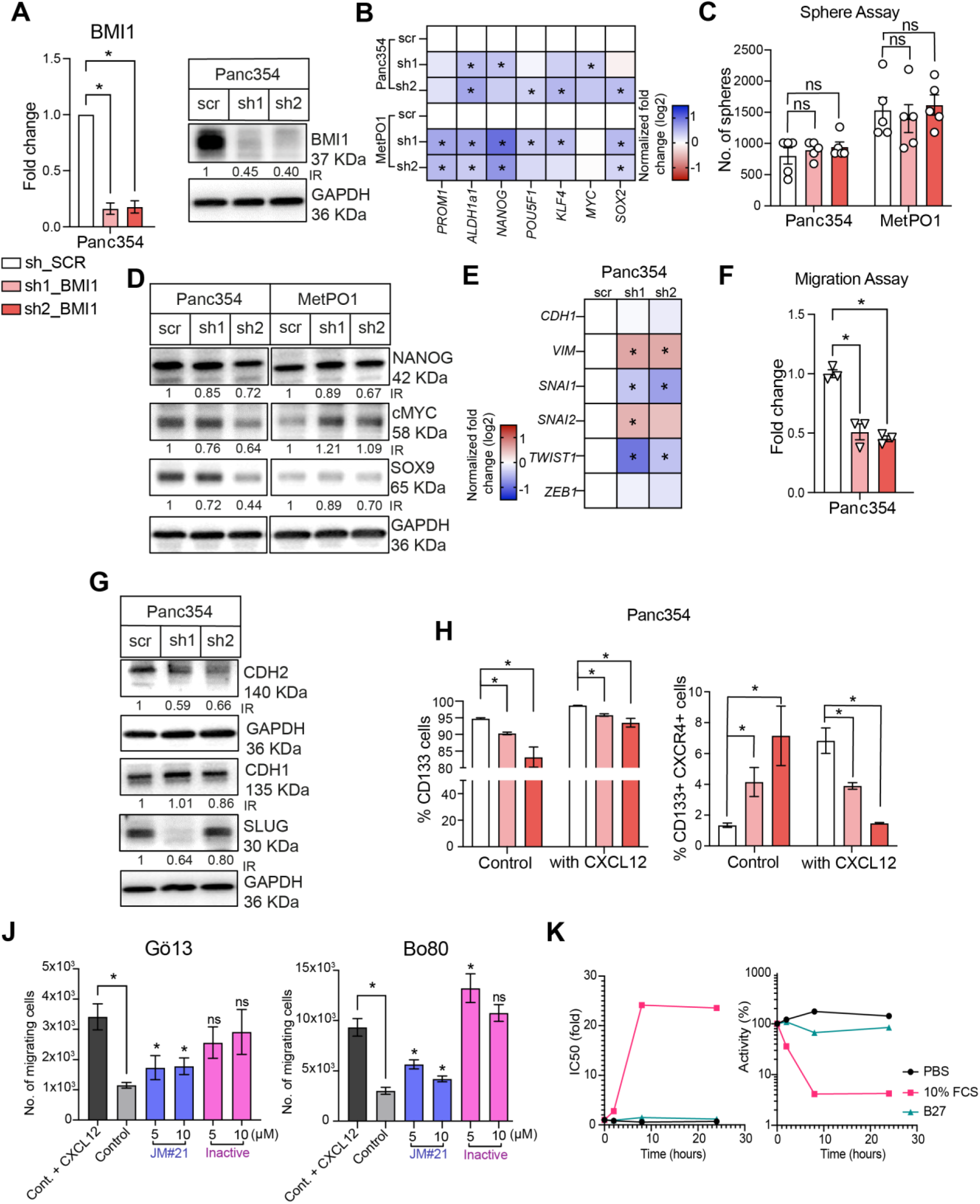
(A) *BMI1* gene expression analysis and western blot analysis in Panc354. GAPDH was used as a loading control. (B) Gene expression analysis for indicated cell lines with genes involved in stemness using qRT-PCR. (C) Sphere formation assays for depicted cell lines. (D) Western blot analysis of NANOG, c-MYC and SOX9 for indicated cell lines. GAPDH was used as a loading control. (E) Gene expression analysis for Panc354 cell line with genes involved in EMT using qRT-PCR. (F) Migration assays towards serum containing media. (G) Western blot analysis of CADHERIN-1, CADHERIN-2 and SLUG for Panc354 cell line. GAPDH was used as a loading control. (H) Flow cytometry analysis of CD133+ cells and CD133+ CXCR4+ cells in Panc354 (sh_*SCR*, sh1_*BMI1* and sh2_*BMI1*) treated with (or without) CXCL12. (J) Migration assays towards CXCL12 for indicated cell lines using JM#21 and the inactive peptide at depicted concentrations. (K) Peptide stability assays for JM#21 in PBS, 10% FCS and B27 to detect activity up to 24 hours. n≥3 for all experiments. *p < 0.05, ns = not significant.

**Supplementary Figure 3.**
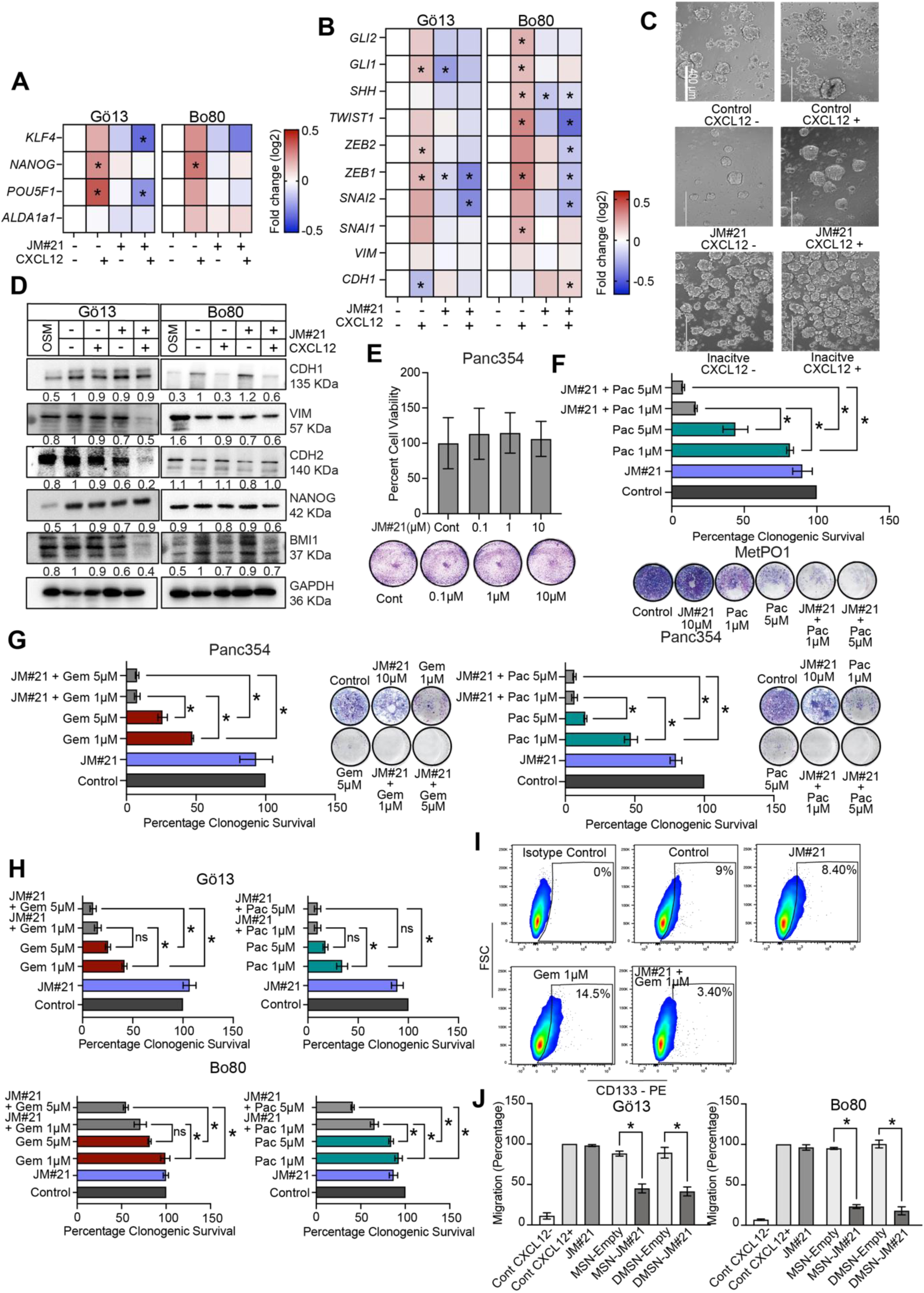
(A) Gene expression analysis for indicated cell lines with genes involved in stemness. (B) Gene expression analysis for indicated cell lines with genes involved in EMT and SHH pathway. (C) Representative micrographs of sphere cultures after 7 days in Panc354 cells after the indicated treatments. JM#21 and inactive peptide pre-treatment was applied for 30 minutes, respectively. (D) Western blot analysis of CADHERIN-1, VIMENTIN, CADHERIN-2, NANOG and BMI1 for indicated cell lines. GAPDH was used as a loading control. (E) Quantification of cell viability and representative pictures for clonogenic assays after 48 hours of JM#21 treatment for CXCL12 pretreated Panc354 cell line. (F) Quantification of cell viability and representative pictures for clonogenic assays after treatment with JM#21 (10*μ*M), gemcitabine (Gem) or paclitaxel (Pac) for indicated concentrations as depicted in experimental design in MetPO1, (G) in Panc354 and (H) Gö13 and Bo80 cell lines. (I) Representative cytometry plots for Panc354 cells. (J) Migration assays towards CXCL12 in FBS-containing medium with the indicated compounds tested in Gö13 and Bo80 cell lines.

## References

Aier, I., Semwal, R., Sharma, A., and Varadwaj, P.K. (2019). A systematic assessment of statistics, risk factors, and underlying features involved in pancreatic cancer. Cancer Epidemiol 58, 104–110.

Alonso-Nocelo, M., Ruiz-Canas, L., Sancho, P., Gorgulu, K., Alcala, S., Pedrero, C., Vallespinos, M., Lopez-Gil, J.C., Ochando, M., Garcia-Garcia, E., et al. (2023). Macrophages direct cancer cells through a LOXL2-mediated metastatic cascade in pancreatic ductal adenocarcinoma. Gut 72, 345–359.

Beitzinger, B., Gerbl, F., Vomhof, T., Schmid, R., Noschka, R., Rodriguez, A., Wiese, S., Weidinger, G., Standker, L., Walther, P., et al. (2021). Delivery by Dendritic Mesoporous Silica Nanoparticles Enhances the Antimicrobial Activity of a Napsin-Derived Peptide Against Intracellular Mycobacterium tuberculosis. Adv Healthc Mater 10, e2100453.

Beitzinger, B., Schmid, R., Jung, C., Tiwary, K., Hermann, P., Jacob, T., and Linden, M. (2024). Confinement and Polarity Effects on the Peptide Packing Density on Mesoporous Silica Nanoparticles. Langmuir 40, 4294–4305.

Bjanes, T.K., Jordheim, L.P., Schjott, J., Kamceva, T., Cros-Perrial, E., Langer, A., Ruiz de Garibay, G., Kotopoulis, S., McCormack, E., and Riedel, B. (2020). Intracellular Cytidine Deaminase Regulates Gemcitabine Metabolism in Pancreatic Cancer Cell Lines. Drug Metab Dispos 48, 153–158.

Bleul, C.C., Farzan, M., Choe, H., Parolin, C., Clark-Lewis, I., Sodroski, J., and Springer, T.A. (1996). The lymphocyte chemoattractant SDF-1 is a ligand for LESTR/fusin and blocks HIV-1 entry. Nature 382, 829–833.

Cash, T.P., Alcala, S., Rico-Ferreira, M.D.R., Hernandez-Encinas, E., Garcia, J., Albarran, M.I., Valle, S., Munoz, J., Martinez-Gonzalez, S., Blanco-Aparicio, C., et al. (2020). Induction of Lysosome Membrane Permeabilization as a Therapeutic Strategy to Target Pancreatic Cancer Stem Cells. Cancers (Basel) 12.

Cespedes, M.V., Unzueta, U., Avino, A., Gallardo, A., Alamo, P., Sala, R., Sanchez-Chardi, A., Casanova, I., Mangues, M.A., Lopez-Pousa, A., et al. (2018). Selective depletion of metastatic stem cells as therapy for human colorectal cancer. EMBO Mol Med 10.

Chen, J., Wang, Z., Gao, S., Wu, K., Bai, F., Zhang, Q., Wang, H., Ye, Q., Xu, F., Sun, H., et al. (2021). Human drug efflux transporter ABCC5 confers acquired resistance to pemetrexed in breast cancer. Cancer Cell Int 21, 136.

Cheung, E.C., DeNicola, G.M., Nixon, C., Blyth, K., Labuschagne, C.F., Tuveson, D.A., and Vousden, K.H. (2020). Dynamic ROS Control by TIGAR Regulates the Initiation and Progression of Pancreatic Cancer. Cancer Cell 37, 168–182 e164.

Erkan, M., Hausmann, S., Michalski, C.W., Fingerle, A.A., Dobritz, M., Kleeff, J., and Friess, H. (2012). The role of stroma in pancreatic cancer: diagnostic and therapeutic implications. Nat Rev Gastroenterol Hepatol 9, 454–467.

Gallmeier, E., Hermann, P.C., Mueller, M.T., Machado, J.G., Ziesch, A., De Toni, E.N., Palagyi, A., Eisen, C., Ellwart, J.W., Rivera, J., et al. (2011). Inhibition of ataxia telangiectasia- and Rad3-related function abrogates the in vitro and in vivo tumorigenicity of human colon cancer cells through depletion of the CD133(+) tumor-initiating cell fraction. Stem Cells 29, 418–429.

Hagmann, W., Faissner, R., Schnolzer, M., Lohr, M., and Jesnowski, R. (2010). Membrane drug transporters and chemoresistance in human pancreatic carcinoma. Cancers (Basel) 3, 106–125.

Harms, M., Habib, M.M.W., Nemska, S., Nicolo, A., Gilg, A., Preising, N., Sokkar, P., Carmignani, S., Raasholm, M., Weidinger, G., et al. (2021). An optimized derivative of an endogenous CXCR4 antagonist prevents atopic dermatitis and airway inflammation. Acta pharmaceutica Sinica B 11, 2694–2708.

Hermann, P.C., Huber, S.L., Herrler, T., Aicher, A., Ellwart, J.W., Guba, M., Bruns, C.J., and Heeschen, C. (2007). Distinct populations of cancer stem cells determine tumor growth and metastatic activity in human pancreatic cancer. Cell Stem Cell 1, 313–323.

Hermann, P.C., Trabulo, S.M., Sainz, B., Jr., Balic, A., Garcia, E., Hahn, S.A., Vandana, M., Sahoo, S.K., Tunici, P., Bakker, A., et al. (2013). Multimodal Treatment Eliminates Cancer Stem Cells and Leads to Long-Term Survival in Primary Human Pancreatic Cancer Tissue Xenografts. PLoS One 8, e66371.

Hu, J.X., Zhao, C.F., Chen, W.B., Liu, Q.C., Li, Q.W., Lin, Y.Y., and Gao, F. (2021). Pancreatic cancer: A review of epidemiology, trend, and risk factors. World J Gastroenterol 27, 4298–4321.

Hu, Q., Qin, Y., Xiang, J., Liu, W., Xu, W., Sun, Q., Ji, S., Liu, J., Zhang, Z., Ni, Q., et al. (2018). dCK negatively regulates the NRF2/ARE axis and ROS production in pancreatic cancer. Cell Prolif 51, e12456.

Jensen, L.J., Kuhn, M., Stark, M., Chaffron, S., Creevey, C., Muller, J., Doerks, T., Julien, P., Roth, A., Simonovic, M., et al. (2009). STRING 8--a global view on proteins and their functional interactions in 630 organisms. Nucleic Acids Res 37, D412–416.

Jiang, L., Wu, J., Yang, Y., Liu, L., Song, L., Li, J., and Li, M. (2012). Bmi-1 promotes the aggressiveness of glioma via activating the NF-kappaB/MMP-9 signaling pathway. BMC Cancer 12, 406.

Kato, T., Matsuo, Y., Ueda, G., Murase, H., Aoyama, Y., Omi, K., Hayashi, Y., Imafuji, H., Saito, K., Morimoto, M., et al. (2022). Enhanced CXCL12/CXCR4 signaling increases tumor progression in radiation-resistant pancreatic cancer. Oncol Rep 47.

Li, C., Heidt, D.G., Dalerba, P., Burant, C.F., Zhang, L., Adsay, V., Wicha, M., Clarke, M.F., and Simeone, D.M. (2007). Identification of pancreatic cancer stem cells. Cancer Res 67, 1030–1037.

Li, C., Wu, J.J., Hynes, M., Dosch, J., Sarkar, B., Welling, T.H., Pasca di Magliano, M., and Simeone, D.M. (2011). c-Met is a marker of pancreatic cancer stem cells and therapeutic target. Gastroenterology 141, 2218–2227 e2215.

Ligorio, M., Sil, S., Malagon-Lopez, J., Nieman, L.T., Misale, S., Di Pilato, M., Ebright, R.Y., Karabacak, M.N., Kulkarni, A.S., Liu, A., et al. (2019). Stromal Microenvironment Shapes the Intratumoral Architecture of Pancreatic Cancer. Cell 178, 160–175 e127.

Liu, S., Dontu, G., Mantle, I.D., Patel, S., Ahn, N.S., Jackson, K.W., Suri, P., and Wicha, M.S. (2006). Hedgehog signaling and Bmi-1 regulate self-renewal of normal and malignant human mammary stem cells. Cancer Res 66, 6063–6071.

Lonardo, E., Frias-Aldeguer, J., Hermann, P.C., and Heeschen, C. (2012). Pancreatic stellate cells form a niche for cancer stem cells and promote their self-renewal and invasiveness. Cell Cycle 11, 1282–1290.

Lonardo, E., Hermann, P.C., Mueller, M.T., Huber, S., Balic, A., Miranda-Lorenzo, I., Zagorac, S., Alcala, S., Rodriguez-Arabaolaza, I., Ramirez, J.C., et al. (2011). Nodal/Activin signaling drives self-renewal and tumorigenicity of pancreatic cancer stem cells and provides a target for combined drug therapy. Cell Stem Cell 9, 433–446.

Lopez-Gil, J.C., Martin-Hijano, L., Hermann, P.C., and Sainz, B., Jr. (2021). The CXCL12 Crossroads in Cancer Stem Cells and Their Niche. Cancers (Basel) 13.

Marechal, R., Demetter, P., Nagy, N., Berton, A., Decaestecker, C., Polus, M., Closset, J., Deviere, J., Salmon, I., and Van Laethem, J.L. (2009). High expression of CXCR4 may predict poor survival in resected pancreatic adenocarcinoma. Br J Cancer 100, 1444–1451.

Mates, J.M., Perez-Gomez, C., and Nunez de Castro, I. (1999). Antioxidant enzymes and human diseases. Clin Biochem 32, 595–603.

Miranda-Lorenzo, I., Dorado, J., Lonardo, E., Alcala, S., Serrano, A.G., Clausell-Tormos, J., Cioffi, M., Megias, D., Zagorac, S., Balic, A., et al. (2014). Intracellular autofluorescence: a biomarker for epithelial cancer stem cells. Nat Methods 11, 1161–1169.

Mirzaei, S., Paskeh, M.D.A., Entezari, M., Mirmazloomi, S.R., Hassanpoor, A., Aboutalebi, M., Rezaei, S., Hejazi, E.S., Kakavand, A., Heidari, H., et al. (2022). SOX2 function in cancers: Association with growth, invasion, stemness and therapy response. Biomedicine & pharmacotherapy = Biomedecine & pharmacotherapie 156, 113860.

Molofsky, A.V., Pardal, R., Iwashita, T., Park, I.K., Clarke, M.F., and Morrison, S.J. (2003). Bmi-1 dependence distinguishes neural stem cell self-renewal from progenitor proliferation. Nature 425, 962–967.

Mueller, M.T., Hermann, P.C., Witthauer, J., Rubio-Viqueira, B., Leicht, S.F., Huber, S., Ellwart, J.W., Mustafa, M., Bartenstein, P., D’Haese, J.G., et al. (2009). Combined targeted treatment to eliminate tumorigenic cancer stem cells in human pancreatic cancer. Gastroenterology 137, 1102–1113.

Muller, A., Homey, B., Soto, H., Ge, N., Catron, D., Buchanan, M.E., McClanahan, T., Murphy, E., Yuan, W., Wagner, S.N., et al. (2001). Involvement of chemokine receptors in breast cancer metastasis. Nature 410, 50–56.

Nguyen, P.H., Giraud, J., Chambonnier, L., Dubus, P., Wittkop, L., Belleannee, G., Collet, D., Soubeyran, I., Evrard, S., Rousseau, B., et al. (2017). Characterization of Biomarkers of Tumorigenic and Chemoresistant Cancer Stem Cells in Human Gastric Carcinoma. Clin Cancer Res 23, 1586–1597.

Oberlin, E., Amara, A., Bachelerie, F., Bessia, C., Virelizier, J.L., Arenzana-Seisdedos, F., Schwartz, O., Heard, J.M., Clark-Lewis, I., Legler, D.F., et al. (1996). The CXC chemokine SDF-1 is the ligand for LESTR/fusin and prevents infection by T-cell-line-adapted HIV-1. Nature 382, 833–835.

Pan, B., Liao, Q., Niu, Z., Zhou, L., and Zhao, Y. (2015). Cancer-associated fibroblasts in pancreatic adenocarcinoma. Future Oncol 11, 2603–2610.

Rahib, L., Smith, B.D., Aizenberg, R., Rosenzweig, A.B., Fleshman, J.M., and Matrisian, L.M. (2014). Projecting cancer incidence and deaths to 2030: the unexpected burden of thyroid, liver, and pancreas cancers in the United States. Cancer Res 74, 2913–2921.

Rasheed, Z.A., Yang, J., Wang, Q., Kowalski, J., Freed, I., Murter, C., Hong, S.M., Koorstra, J.B., Rajeshkumar, N.V., He, X., et al. (2010). Prognostic significance of tumorigenic cells with mesenchymal features in pancreatic adenocarcinoma. J Natl Cancer Inst 102, 340–351.

Rosenholm, J.M., Meinander, A., Peuhu, E., Niemi, R., Eriksson, J.E., Sahlgren, C., and Linden, M. (2009). Targeting of porous hybrid silica nanoparticles to cancer cells. ACS Nano 3, 197–206.

Sainz, B., Jr., Alcala, S., Garcia, E., Sanchez-Ripoll, Y., Azevedo, M.M., Cioffi, M., Tatari, M., Miranda-Lorenzo, I., Hidalgo, M., Gomez-Lopez, G., et al. (2015). Microenvironmental hCAP-18/LL-37 promotes pancreatic ductal adenocarcinoma by activating its cancer stem cell compartment. Gut 64, 1921–1935.

Seemann, S., and Lupp, A. (2015). Administration of a CXCL12 Analog in Endotoxemia Is Associated with Anti-Inflammatory, Anti-Oxidative and Cytoprotective Effects In Vivo. PLoS One 10, e0138389.

Shen, B., Zheng, M.Q., Lu, J.W., Jiang, Q., Wang, T.H., and Huang, X.E. (2013). CXCL12-CXCR4 promotes proliferation and invasion of pancreatic cancer cells. Asian Pac J Cancer Prev 14, 5403–5408.

Smigiel, J.M., Parameswaran, N., and Jackson, M.W. (2017). Potent EMT and CSC Phenotypes Are Induced By Oncostatin-M in Pancreatic Cancer. Mol Cancer Res 15, 478–488.

Song, L.B., Li, J., Liao, W.T., Feng, Y., Yu, C.P., Hu, L.J., Kong, Q.L., Xu, L.H., Zhang, X., Liu, W.L., et al. (2009). The polycomb group protein Bmi-1 represses the tumor suppressor PTEN and induces epithelial-mesenchymal transition in human nasopharyngeal epithelial cells. J Clin Invest 119, 3626–3636.

Sperti, C., Pasquali, C., Piccoli, A., and Pedrazzoli, S. (1997). Recurrence after resection for ductal adenocarcinoma of the pancreas. World J Surg 21, 195–200.

Tang, Z., Li, C., Kang, B., Gao, G., Li, C., and Zhang, Z. (2017). GEPIA: a web server for cancer and normal gene expression profiling and interactive analyses. Nucleic Acids Res 45, W98–W102.

Thomas, S.K., Lee, J., and Beatty, G.L. (2020). Paracrine and cell autonomous signalling in pancreatic cancer progression and metastasis. EBioMedicine 53, 102662.

Tu, Z., Xie, S., Xiong, M., Liu, Y., Yang, X., Tembo, K.M., Huang, J., Hu, W., Huang, X., Pan, S., et al. (2017). CXCR4 is involved in CD133-induced EMT in non-small cell lung cancer. Int J Oncol 50, 505–514.

Valk-Lingbeek, M.E., Bruggeman, S.W., and van Lohuizen, M. (2004). Stem cells and cancer; the polycomb connection. Cell 118, 409–418.

Vora, P., Seyfrid, M., Venugopal, C., Qazi, M.A., Salim, S., Isserlin, R., Subapanditha, M., O’Farrell, E., Mahendram, S., Singh, M., et al. (2019). Bmi1 regulates human glioblastoma stem cells through activation of differential gene networks in CD133+ brain tumor initiating cells. J Neurooncol 143, 417–428.

Walter, K., Tiwary, K., Trajkovic-Arsic, M., Hidalgo-Sastre, A., Dierichs, L., Liffers, S.T., Gu, J., Gout, J., Schulte, L.A., Munch, J., et al. (2019). MEK Inhibition Targets Cancer Stem Cells and Impedes Migration of Pancreatic Cancer Cells In Vitro and In Vivo. Stem Cells Int 2019, 8475389.

Widschwendter, M., Fiegl, H., Egle, D., Mueller-Holzner, E., Spizzo, G., Marth, C., Weisenberger, D.J., Campan, M., Young, J., Jacobs, I., et al. (2007). Epigenetic stem cell signature in cancer. Nat Genet 39, 157–158.

Xu, X., Su, B., Xie, C., Wei, S., Zhou, Y., Liu, H., Dai, W., Cheng, P., Wang, F., Xu, X., et al. (2014). Sonic hedgehog-Gli1 signaling pathway regulates the epithelial mesenchymal transition (EMT) by mediating a new target gene, S100A4, in pancreatic cancer cells. PLoS One *9*, e96441.

Xu, X., Zhou, Y., Xie, C., Wei, S.M., Gan, H., He, S., Wang, F., Xu, L., Lu, J., Dai, W., et al. (2012). Genome-wide screening reveals an EMT molecular network mediated by Sonic hedgehog-Gli1 signaling in pancreatic cancer cells. PLoS One 7, e43119.

Yin, T., Zhang, Z., Cao, B., Duan, Q., Shi, P., Zhao, H., Camara, S.N., Shen, Q., and Wang, C. (2016). Bmi1 inhibition enhances the sensitivity of pancreatic cancer cells to gemcitabine. Oncotarget 7, 37192–37204.

Zhang, J., Liu, C., Mo, X., Shi, H., and Li, S. (2018). Mechanisms by which CXCR4/CXCL12 cause metastatic behavior in pancreatic cancer. Oncol Lett 15, 1771–1776.

Zhang, L., Li, J., Zong, L., Chen, X., Chen, K., Jiang, Z., Nan, L., Li, X., Li, W., Shan, T., et al. (2016). Reactive Oxygen Species and Targeted Therapy for Pancreatic Cancer. Oxid Med Cell Longev 2016, 1616781.

Zhang, N.H., Li, J., Li, Y., Zhang, X.T., Liao, W.T., Zhang, J.Y., Li, R., and Luo, R.C. (2012). Co-expression of CXCR4 and CD133 proteins is associated with poor prognosis in stage II-III colon cancer patients. Exp Ther Med 3, 973–982.

Zirafi, O., Hermann, P.C., and Munch, J. (2016). Proteolytic processing of human serum albumin generates EPI-X4, an endogenous antagonist of CXCR4. J Leukoc Biol 99, 863–868.

Zirafi, O., Kim, K.A., Standker, L., Mohr, K.B., Sauter, D., Heigele, A., Kluge, S.F., Wiercinska, E., Chudziak, D., Richter, R., et al. (2015). Discovery and characterization of an endogenous CXCR4 antagonist. Cell Rep 11, 737–747.

